# Longitudinal analysis of invariant natural killer T cell activation reveals a cMAF-associated transcriptional state of NKT10 cells

**DOI:** 10.1101/2021.12.29.474454

**Authors:** Harry Kane, Nelson M. LaMarche, Áine Ní Scannail, Michael B. Brenner, Lydia Lynch

## Abstract

Innate T cells, including CD1d-restricted invariant natural killer T (iNKT) cells, are characterized by their rapid activation in response to non-peptide antigens, such as lipids. While the transcriptional profiles of naive, effector and memory adaptive T cells have been well studied, less is known about transcriptional regulation of different iNKT cell activation states. Here, using single cell RNA-sequencing, we performed longitudinal profiling of activated iNKT cells, generating a transcriptomic atlas of iNKT cell activation states. We found that transcriptional signatures of activation are highly conserved among heterogeneous iNKT cell populations, including NKT1, NKT2 and NKT17 subsets, and human iNKT cells. Strikingly, we found that regulatory iNKT cells, such as adipose iNKT cells, undergo blunted activation, and display constitutive enrichment of memory-like cMAF^+^ and KLRG1^+^ populations. Moreover, we identify a conserved cMAF-associated transcriptional network among NKT10 cells, providing novel insights into the biology of regulatory and antigen experienced iNKT cells.

## Introduction

Activation of T cells following recognition of cognate antigen is essential for mounting effective immune responses against pathogens and tumors^1^. Typically, in the case of MHC-restricted adaptive CD4^+^ and CD8^+^ T cells, this requires extensive transcriptional remodeling over several days to facilitate proliferation and differentiation of naive T cells into clonal effector populations that traffic to sites of infection or tissue damage^2^. Transcriptional and metabolic remodeling is also needed to generate memory T cells that can be rapidly reactivated following secondary antigen encounter during reinfection^2^. Innate T cells, including CD1d-restricted invariant natural killer T (iNKT) cells, contrast and complement this paradigm by exiting thymic development as poised ‘effector-memory-like’ cells already capable of mounting potent cytokine responses within minutes of activation. This allows iNKT cells to rapidly transactivate other immune populations and orchestrate immune responses^3,4^. Activation also induces iNKT cell proliferation, generating an expanded pool of effector cells within 72 hours, most of which subsequently undergo apoptosis as the expanded iNKT cell pool contracts within 7 days^5–7^. However, some iNKT cells persist after the immune response subsides^6–8^, and there is evidence that antigen challenge induces long term changes in the iNKT cell repertoire analogous to memory T cell differentiation. For example, several studies have demonstrated that activation of iNKT cells with α-galalctosylceramide (αGalCer), a potent glycolipid antigen, induces the emergence of novel KLRG1^+^ and Follicular Helper iNKT (NKT_FH_) cell populations that are greatly enriched after 3-7 days, and still detectable >30 days after αGalCer challenge^8–11^. However, our knowledge of the transcriptional programs underpinning iNKT cell activation remains limited, and there are also relatively few transcriptional resources available for studying activated iNKT cells, especially compared to adaptive T cells^12^.

Analysis and interpretation of iNKT cell biology is also challenging because iNKT cells exhibit heterogeneity, including NKT1, NKT2 and NKT17 subsets that broadly mirror CD4^+^ Th1, Th2 and Th17 cells^13^. Past studies of NKT1, NKT2 and NKT17 subsets largely focused on iNKT cell thymic development or steady state phenotype in the absence of activation^13–16^, and less is known about iNKT cell subsets after activation. Using parabiosis models, we and others have also shown that iNKT cells are predominantly tissue resident^17^, and that this can strongly influence their biology^18^. For example, iNKT cells resident in adipose tissue exhibit an unusual regulatory phenotype characterized by increased KLRG1 expression, reduced expression of the transcription factor promyelocytic leukemia zinc-finger (PLZF), and increased production of IL-10 through an IRE1a-XBP1s-E4BP4 axis, enabling these cells to suppress inflammation and promote metabolic homeostasis^17,18^. Interestingly, Sag *et al*. (2014) demonstrated that IL-10^+^ iNKT (NKT10) cells emerge in other organs such as the spleen after repeated antigen challenge^19^, indicating that TCR stimulation can induce a regulatory phenotype, and that NKT10 cells can potentially be considered a memory-like population. However, the relationship between NKT10 cells and other memory-like populations, such as KLRG1^+^ and NKT_FH_ cells, remains unclear. Furthermore, it is unknown whether similar factors regulate NKT10 cells present in adipose tissue versus those induced after antigen challenge.

To characterize transcriptional remodeling in activated iNKT cells while also considering subset and tissue-associated heterogeneity, we performed single cell RNA-Sequencing (scRNA-Seq) of 48,813 murine iNKT cells from spleen and adipose tissue at steady state and 4 hours, 72 hours and 4 weeks after *in vivo* stimulation with αGalCer, as well as after repeated αGalCer challenge. We also reanalyzed published human and murine data to generate a transcriptomic atlas of iNKT cell activation states. We found that activation induces rapid and extensive transcriptional remodeling in iNKT cells, and that a common transcriptional framework underpins the activation of diverse iNKT cell populations. However, regulatory iNKT cell populations demonstrate largely blunted activation in response to αGalCer, and display enrichment of memory-like KLRG1^+^ and cMAF^+^ iNKT cell subsets expressing a T regulatory type 1 (Tr1) cell gene signature. We also show that cMAF^+^ iNKT cells are enriched for NKT10 cells, and express a gene signature similar to NKT_FH_ cells. Overall, this study provides novel insights into longitudinal transcriptional remodeling in activated iNKT cells and the phenotype of regulatory iNKT cells, while also generating a novel transcriptomic resource for interrogation of iNKT cell biology.

## Results

### iNKT cells undergo rapid and extensive transcriptional remodeling in response to αGalCer

To investigate transcriptional remodeling in activated iNKT cells we performed 10x scRNA-Seq of whole murine adipose and splenic iNKT cells 4 hours, 72 hours and 4 weeks after *in vivo* stimulation with αGalCer, and reanalyzed our published scRNA-Seq of steady state murine adipose and splenic iNKT cells^18^ (GSE142845, Figure 1A). We first analyzed our steady state, 4 hour and 72 hour splenic iNKT cell data. After quality control measures we obtained 16,701 splenic iNKT cells, including >4000 cells per activation state. After performing UMAP we observed minimal overlap between iNKT cells from different activation states (Figure 1B), indicating that iNKT cells undergo rapid and extensive transcriptional remodeling during early activation. Using gene expression analysis (Table S1) we found that steady state iNKT cells displayed enrichment of NKT1 and NKT17 cell markers such as *Il2rb*, *Klrb1c*, *Rorc*, and *Il7r* (Figure 1C)^13^, but following activation iNKT cells rapidly downregulated these genes within 4 hours, and upregulated expression of T cell activation markers and cytokines, including *Il2ra*, *Irf4*, *Nr4a1*, *Pdcd1*, *Ifng*, *Il4* and *Il17a* (Figure 1C). This was accompanied by increased expression of *Zbtb16* (PLZF) and the PLZF regulon genes *Icos* and *Cd40lg* (Figure 1C), consistent with published data demonstrating that PLZF is required for the innate response of iNKT cells to antigen^20^. Activated iNKT cells also downregulated expression of the transcription factor *Id2* (Figure 1C), which plays an essential role in normal iNKT cell activation^21^, and upregulated expression of genes regulating T cell metabolic activation, including *Myc*, *Hif1a* and *Tfrc*^22–24^, suggesting that activated iNKT cells undergo metabolic remodeling.

**Figure 1:**
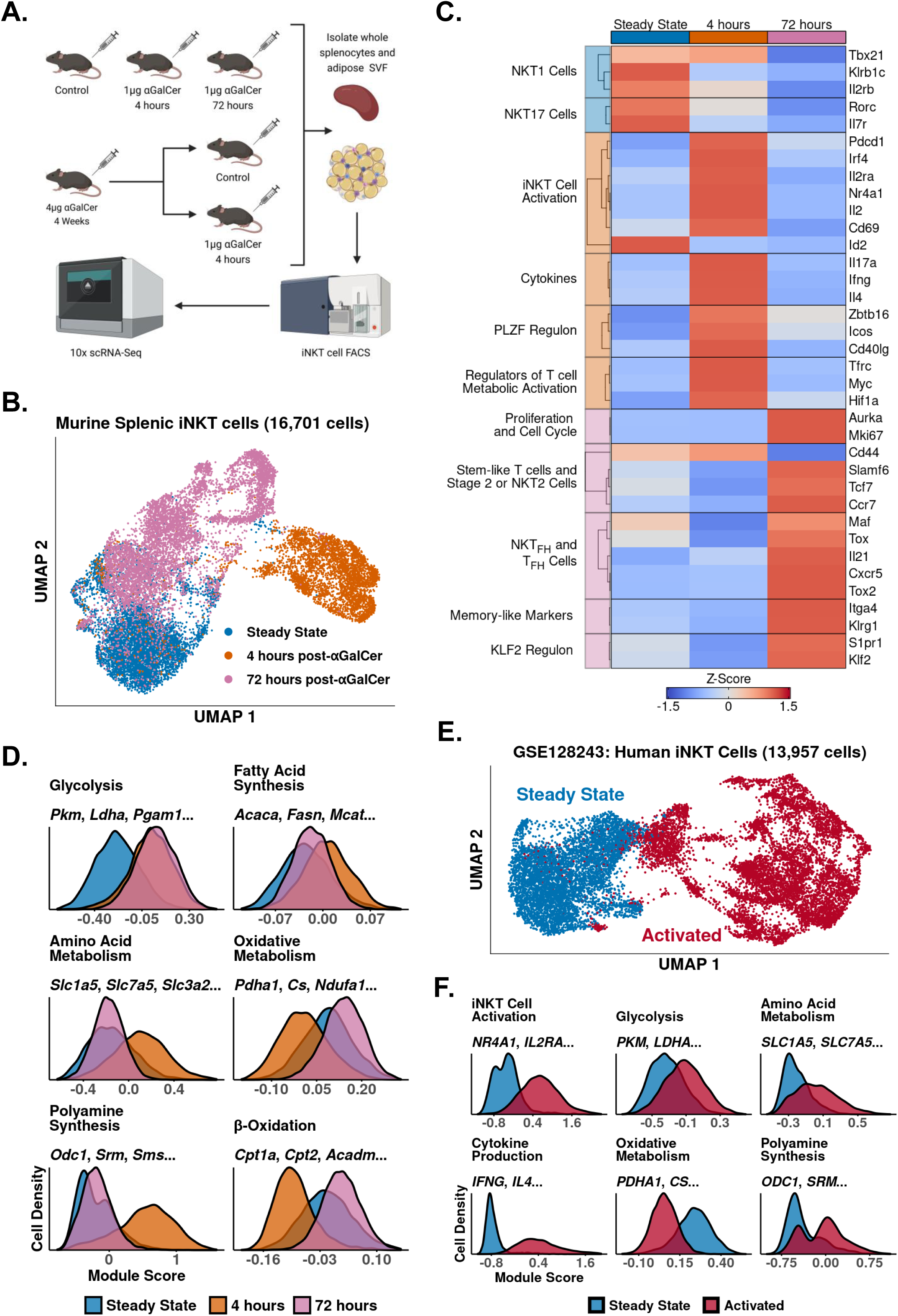
iNKT cells undergo rapid and extensive transcriptional remodeling in response to αGalCer. **A.** Cartoon illustrating the experimental design for the generation of all scRNA-Seq data. **B.** UMAP of murine splenic iNKT cells with cell cycle regression. **C.** Heatmap of averaged gene expression with hierarchical clustering in the data from Figure 1B. **D.** Histograms showing expression of metabolic module scores in the data from Figure 1B. **E.** UMAP of human PBMC iNKT cells reanalyzed from GSE128243. **F.** Histograms showing functional and metabolic module scores in the data from Figure 1E.

By 72 hours, however, expression of activation and cytokine genes was greatly reduced, and we identified enrichment of genes associated with proliferation, stem-like T cells, and NKT2 or Stage 2 iNKT cells, including *Mki67*, *Slamf6*, *Tcf7* and *Ccr7* (Figure 1C)^25,26^. We observed that some 72 hour cells displayed enrichment of T_FH_ and NKT_FH_ markers, including *Cxcr5*, *Il21* and *Maf*^9,11,12,27^, and memory-like iNKT cell markers, such as *Itga4* and *Klrg1*^8^ (Figure 1C, Figure S1), corresponding with previous studies documenting the appearance of NKT_FH_ and KLRG1^+^ iNKT cells after αGalCer challenge^8–10,28^. We also found increased expression of genes associated with the KLF2 regulon, including *Klf2* and *S1pr1* (Figure 1C). KLF2 is known to induce T cell thymic egress and trafficking through secondary lymphoid organs^29^; while iNKT cells are generally tissue resident under steady state conditions^17^, expression of *Klf2* and *S1pr1* 72 hours post-αGalCer suggests that activated iNKT cells may traffic to other sites.

Having identified enrichment of *Myc*, *Hif1a* and *Tfrc* 4 hours post-αGalCer, we wondered what type of metabolic remodeling activated iNKT cells undergo *in vivo*. To map metabolic changes during iNKT cell activation we generated gene module scores using the KEGG pathway^30^ and Gene Ontology Consortium^31^ databases (Table S2) and scored our data. We found 4 hour activated cells upregulated glycolysis, amino acid metabolism, polyamine synthesis and fatty acid synthesis signatures, whereas oxidative signatures were downregulated compared to steady state iNKT cells (Figure 1D). This suggests that, despite being poised at steady state for cytokine production, activated iNKT cells, like adaptive T cells, switch on aerobic glycolysis and upregulate biosynthethic pathways to fuel cytokine production, growth and proliferation^22,32,33^. Our data is also consistent with recent work identifying glucose as an important fuel for iNKT cell effector function^34,35^. Interestingly, we found that 72 hour activated cells engage oxidative signatures while maintaining elevated expression of glycolytic genes (Figure 1D), indicating that the metabolic requirements of iNKT cells change across different activation states. We also observed reduced expression of polyamine synthesis and amino acid metabolism signatures 72 hours post-αGalCer, suggesting that those pathways are coupled to early iNKT cell activation and cytokine production, while oxidative metabolism may be coupled to proliferation when the iNKT cell pool expands 4-10 fold *in vivo* by 72 hours^6,7^.

Having profiled transcriptional remodeling in activated murine iNKT cells, we wondered whether similar remodeling occurs in human iNKT cells. To investigate human iNKT cell activation we reanalyzed published scRNA-Seq data of human iNKT cells isolated from peripheral blood mononuclear cells (PBMCs) and stimulated *ex vivo* with phorbol 12-myristate 13-acetate (PMA) and Ionomycin (GSE128243)^36^. Following quality control measures we obtained 13,957 cells, and we found that human iNKT cells also undergo rapid and extensive transcriptional remodeling after activation (Figure 1E, Figure 1F). Furthermore, activated human iNKT cells recapitulated the metabolic reprogramming observed in activated murine iNKT cells, displaying upregulated glycolytic, amino acid metabolism and polyamine synthesis signatures, and reduced expression of oxidative signatures (Figure 1F). Thus, transcriptional signatures of iNKT cell activation are conserved across species.

### Oxidative Phosphorylation differentiates functional responses to αGalCer in NKT2 and NKT17 cells versus NKT1 cells

We next asked whether iNKT cell subsets expressed different transcriptional signatures after activation. We performed subclustering of murine splenic iNKT cells at steady state and 4 hours post-αGalCer, and identified clusters corresponding to NKT1, NKT2 and NKT17 cells (Figure 2A) using the published marker genes *Tbx21*, *Zbtb16*, *Rorc, Ifng*, *Il4* and *Il17a*^5,13^ (Figure 2B). Notably, we found that all subsets upregulated *Zbtb16* and *Tbx21* after activation (Figure 2B), suggesting that PLZF and T-bet play subset-independent roles during activation. When we performed gene expression analysis we found that all subsets demonstrated upregulation of activation, cytokine, glycolysis, amino acid metabolism, polyamine synthesis and fatty acid synthesis signatures after αGalCer (Figure 2C, Supplemental Figure 2), indicating that a common transcriptional framework underpins the activation of functionally diverse iNKT cell subsets. We also identified genes specifically enriched in one or more subsets, such as *Gzmb* and *Ccl4* in NKT1 cells (Figure 2C; Table S3). Strikingly, we found that NKT2 and NKT17 cells, but not NKT1 cells, shared expression of many genes, including *Lif*, *Lta*, *Cd274* and *Ncoa7* (Figure 2C). Activated NKT2 and NKT17 cells also demonstrated increased whole transcriptome correlation compared to activated NKT1 cells (Figure 2D), indicating that activated NKT2 and NKT17 cells are transcriptionally similar compared to NKT1 cells.

**Figure 2:**
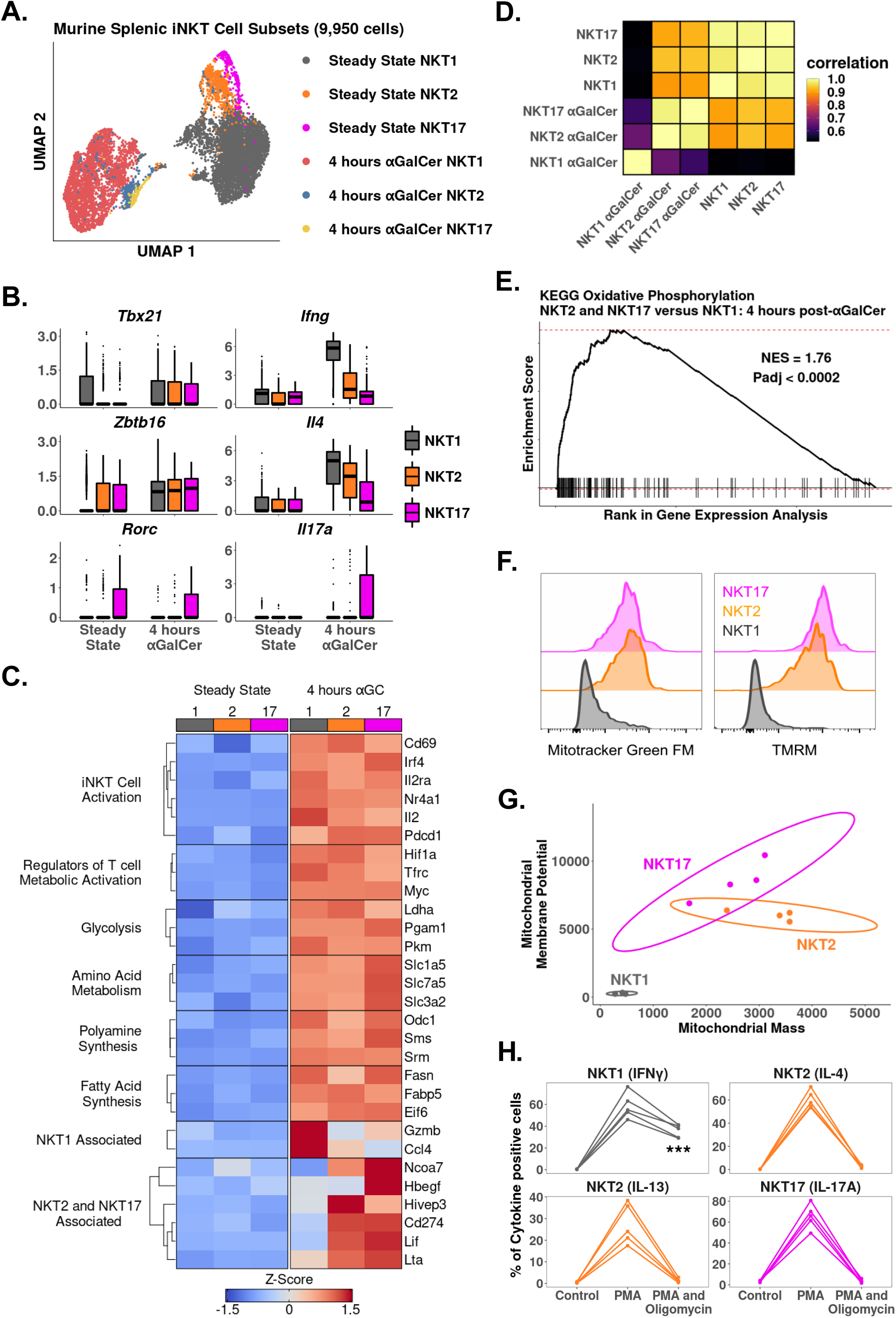
Oxidative Phosphorylation differentiates functional responses to αGalCer in NKT2 and NKT17 cells versus NKT1 cells. **A.** UMAP of murine splenic iNKT cells. **B.** Boxplots showing selected gene expression in the data from Figure 2A. **C.** Heatmap of averaged gene expression with hierarchical clustering in NKT1 (1), NKT2 (2) and NKT17 cells (17) from the data in Figure 2A. **D.** Correlation plot of total normalized RNA counts for all genes across NKT1, NKT2 and NKT17 cell subsets at steady state or 4 hours post-αGalCer. **E.** GSEA plot of KEGG Oxidative Phosphorylation comparing NKT2 cells and NKT17 cells versus NKT1 cells 4 hours post-αGalCer. “NES” denotes Normalized Enrichment Score and “Padj” denotes the adjusted P value for the enrichment. **F.** Histograms showing staining of Mitotracker Green FM and TMRM in CD44^+^ NKT1, NKT2 and NKT17 cells from BALB/C mouse thymus. Histograms were normalized to the mode. iNKT cells were defined as live, single CD19^-^ CD8^-^ CD3^low^ CD1d Tetramer^+^ cells. **G.** Scatter plot showing median fluorescence intensity (MFI) of N = 4 biological replicates from one experiment. Experiment performed at least twice. Ellipses denote 95% confidence intervals of the replicate distributions for each iNKT cell subset. **H.** Line plots showing production of flagship cytokines by CD44^+^ BALB/C mouse thymic iNKT cells: NKT1 (IFNγ), NKT2 (IL-4 and IL-13), and NKT17 cells (IL-17A) without stimulation, after 50ng PMA and 1µg Ionomycin, and after 50ng PMA and 1µg Ionomycin in the presence of 40nM Oligomycin (all 4 hours *ex vivo*). N = 5 biological replicates from one experiment. Experiment performed at least twice. One way ANOVA and Tukey’s post hoc test. Asterisks denote significance, * padj < 0.05; ** padj <0.01; *** padj < 0.001.

To investigate the shared transcriptional signatures of NKT2 and NKT17 cells, we performed GSEA^37,38^ comparing activated NKT2 and NKT17 cells versus activated NKT1 cells using the KEGG pathway database^30^. We identified enrichment of Oxidative Phosphorylation (Figure 2E), suggesting that NKT2 and NKT17 cells use oxidative metabolism more than NKT1 cells. To validate this result we first measured mitochondrial mass and membrane potential in thymic CD44^+^ NKT1, NKT2 and NKT17 cells, and we found that NKT2 and NKT17 cells had significantly increased mitochondrial mass and membrane potential compared to NKT1 cells (Figure 2F and 2G). We next investigated whether NKT2 and NKT17 cells were more functionally dependent on oxidative metabolism than NKT1 cells by stimulating iNKT cells *ex vivo* with PMA and Ionomycin for 4 hours in the presence or absence of oligomycin, to inhibit oxidative phosphorylation^39^. Treatment with oligomycin globally reduced cytokine production across all iNKT cell subsets, however, we found that production of IL-4, IL-13 and IL-17A was almost completely ablated compared to production of IFNγ (Figure 2H), demonstrating that oxidative metabolism is essential for NKT2 and NKT17 cytokines but less so for NKT1 cytokines.

### Adipose iNKT cells display blunted and delayed activation after αGalCer, enrichment of Tr1 Cell markers, and hallmarks of chronic endogenous activation

We and others have shown that adipose iNKT cells display an unusual regulatory phenotype characterized by E4BP4 (*Nfil3*) expression and enrichment of NKT10 cells^17–19^. Having characterized activated splenic iNKT cells, we next investigated activated adipose iNKT cells. Combined analysis of adipose and splenic iNKT cells at steady state, 4 hours post-αGalCer, and 72 hours post-αGalCer returned 28,561 cells, including 11,860 adipose iNKT cells, and >2900 cells per activation state. When we performed UMAP we found that adipose and splenic iNKT cells displayed minimal overlap (Figure 3A), indicative of constitutive transcriptional differences between adipose and splenic iNKT cells. Gene expression analysis identified 971 genes enriched among all adipose iNKT cells regardless of activation status or subset, versus only 65 genes enriched among all splenic iNKT cells (Table S4), indicating that adipose but not splenic iNKT cells retain expression of many conserved genes during activation. When we profiled these conserved genes using over-representation analysis against the KEGG pathway database we identified Ribosome as the sole enriched pathway among splenic iNKT cells, whereas adipose iNKT cells displayed enrichment of pathways related to uptake from the extracellular environment (Endocytosis, Regulation of actin cytoskeleton, FcγR-mediated phagocytosis), adhesion (Focal adhesion, Leukocyte transendothelial migration), cytotoxicity and cell death (NK cell mediated cytotoxicity, Apoptosis), cellular senescence, chemokine signaling, and TCR signaling; (Figure 3B).

**Figure 3:**
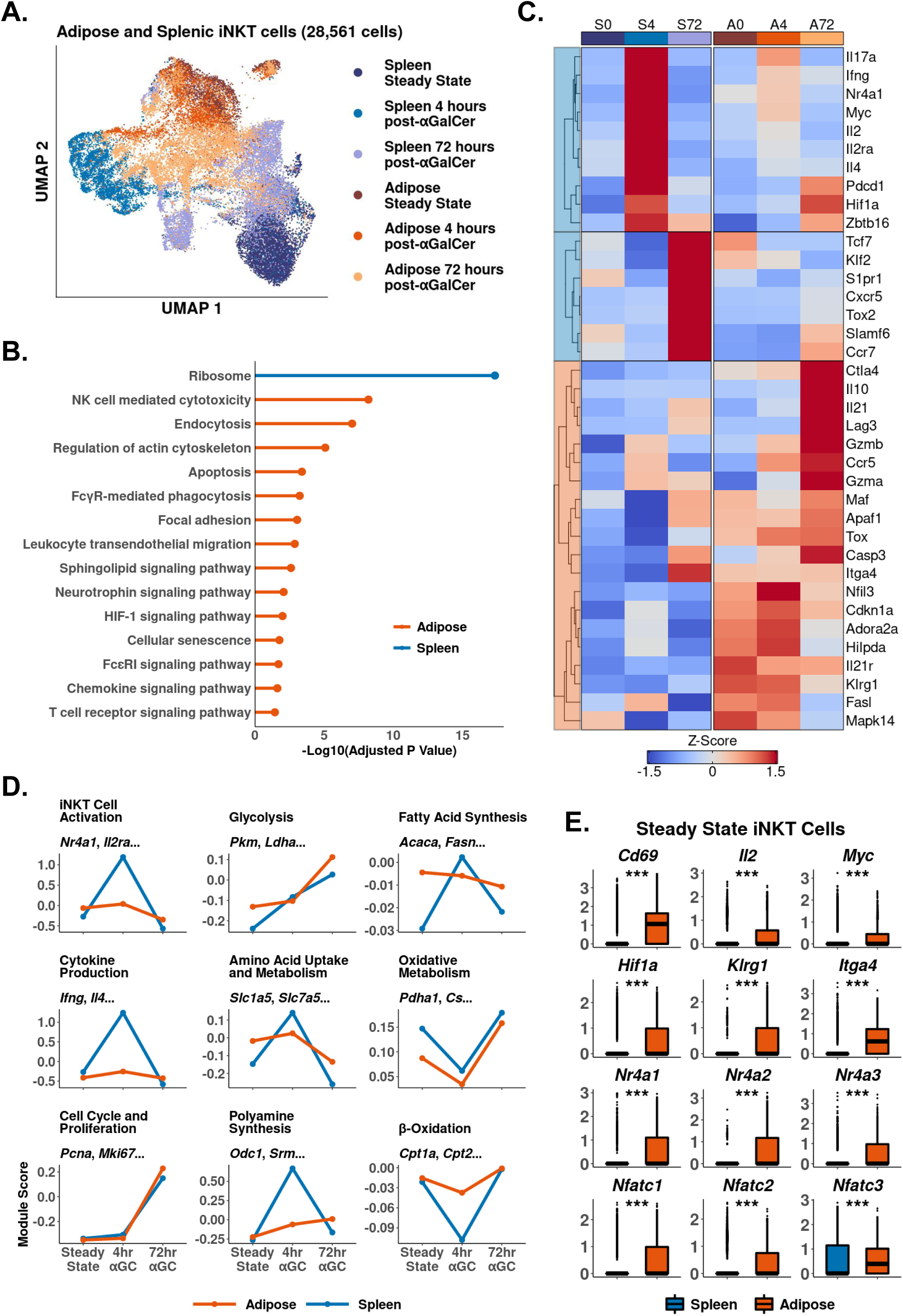
Adipose iNKT cells display blunted and delayed activation after αGalCer, enrichment of Tr1 Cell markers, and hallmarks of chronic endogenous activation. **A**. UMAP of murine adipose and splenic iNKT cells with cell cycle regression. **B.** Lollipop plot showing enrichment of non-disease KEGG pathways among 972 genes conserved in adipose iNKT cells and 65 genes conserved in splenic iNKT cells. **C.** Heatmap of averaged gene expression with hierarchical clustering in the data from Figure 3A. **D.** Line plots showing median expression of functional and metabolic module scores in the data from Figure 4A. **E.** Boxplots showing gene expression in murine steady state splenic or adipose iNKT cells. Horizontal lines represent median values.

We next compared gene expression among individual iNKT cell activation states. Intriguingly, there was greatly reduced expression of activation markers such as *Il2ra* and *Nr4a1* (Nur77) in adipose versus splenic iNKT cells at 4 hours post-αGalCer (Figure 3C). We previously found that Nur77 is enriched in adipose iNKT cells compared to splenic iNKT cells at steady state^17^, but Nur77 expression does not increase upon activation in adipose iNKT cells, unlike splenic iNKT cells. Using module scoring we found that adipose iNKT cells showed a blunted initial response to αGalCer characterized by reduced upregulation of activation and cytokine signatures, and reduced metabolic remodeling (Figure 3D). At 72 hours post-αGalCer, when proliferative gene signatures are upregulated, adipose and splenic iNKT cells showed similar enrichment of proliferation markers (Figure 3D), but adipose iNKT cells had reduced expression of stem-like markers such as *Tcf7* and *Slamf6*, and reduced expression of T_FH_ markers such as *Cxcr5* and *Tox2* (Figure 3C). Adipose iNKT cells showed increased expression of *Il10*, *Lag3*, *Ctla4*, *Il21*, *Ccr5*, *Hif1a*, *Maf*, and *Gzmb* (Figure 3C), which are markers typically associated with Tr1 cells, a heterogeneous population of regulatory T cells that do not express FOXP3^40–44^. We found that adipose iNKT cells do not express FOXP3 and instead express the transcription factor E4BP4 for IL-10 production^17^. Other genes enriched among adipose iNKT cells included *Il21r*, the adenosine receptor *Adora2a*, and the exhaustion marker *Tox* (Figure 3C). In summary, activation with αGalCer induces differential transcriptional remodeling in adipose versus splenic iNKT cells, and the peak of the regulatory response in adipose iNKT cells is delayed compared to the rapid cytokine burst in splenic iNKT cells.

Notably, although adipose iNKT cells displayed blunted activation and metabolic remodeling compared to splenic iNKT cells, adipose iNKT cells were already enriched for activation, glycolysis and amino acid metabolism gene signatures at steady state (Figure 3D), suggesting that adipose cells are activated at baseline. At steady state, adipose iNKT cells had increased expression of key activation markers such as *Il2*, *Cd69*, *Myc*, and *Nr4a* genes, as well as increased *Nfat* gene expression (Figure 3E). *Nr4a* and *Nfat* genes are commonly upregulated in chronically activated T cells, often in combination with *Tox*^45–47^, suggesting that adipose iNKT cells experience chronic activation. Furthermore, we found that steady state adipose iNKT cells demonstrated increased expression of markers of antigen experience, *Klrg1* and *Itga4* (Figure 3E)^8^, and we have previously shown that KLRG1^+^ iNKT cells are greatly enriched in adipose tissue^17^. This indicates that adipose iNKT cells experience chronic endogenous activation, which may dampen their ability to undergo further activation in response to αGalCer, and could explain the nature of their regulatory phenotype.

### scRNA-Seq identifies transcriptional signatures of adipose iNKT cell subset activation and adipose NKT10 cells

We have recently shown that adipose iNKT cells are more heterogeneous than originally anticipated. At steady state, these cells are comprised of NK1.1^+^ NKT1 cells, NK1.1^-^ NKT1 cells, and NKT17 cells^18^. NK1.1^+^ and NK1.1^-^ adipose iNKT cells are functionally distinct, as NK1.1^+^ cells produce IFNγ, and NK1.1^-^ iNKT cells produce IL-4 and IL-10. However, it is unknown if these populations engage distinct molecular programs in response to αGalCer, a proposed therapy for type 2 diabetes and obesity. We also wondered whether the reduced responsiveness to αGalCer in adipose iNKT cells was linked to any particular adipose iNKT cell population. To answer these questions we performed analysis of adipose iNKT cells at steady state and 4 hours post-αGalCer. Unlike splenic iNKT cells, where activation accounted for most of the variance (Figure 2A), transcriptional differences between adipose NKT1 cells and NKT17 cells accounted for most of the variance among adipose iNKT cells (Figure 4A). We found that adipose NKT1 and NKT17 cells both upregulated expression of activation, cytokine and metabolic gene signatures after αGalCer (Figure 4B). Comparison of adipose and splenic iNKT cell subsets showed that all adipose iNKT cell subsets had reduced activation and cytokine gene signature expression, and reduced metabolic remodeling 4 hours post-αGalCer (Figure 4C). This indicates that all adipose iNKT cell subsets respond to αGalCer but no single subset (e.g. NKT1 or NKT17) was uniquely hyporesponsive *vis a vis* the spleen. Interestingly, we identified increased oxidative gene expression in adipose NKT17 cells versus adipose NKT1 cells (Figure 4C), similar to our finding in splenic iNKT cells (Figure 2). We have previously shown that γδ17 cells also display enrichment of oxidative metabolism^39^, suggesting that this is a conserved feature of innate T cells that produce IL-17.

**Figure 4.**
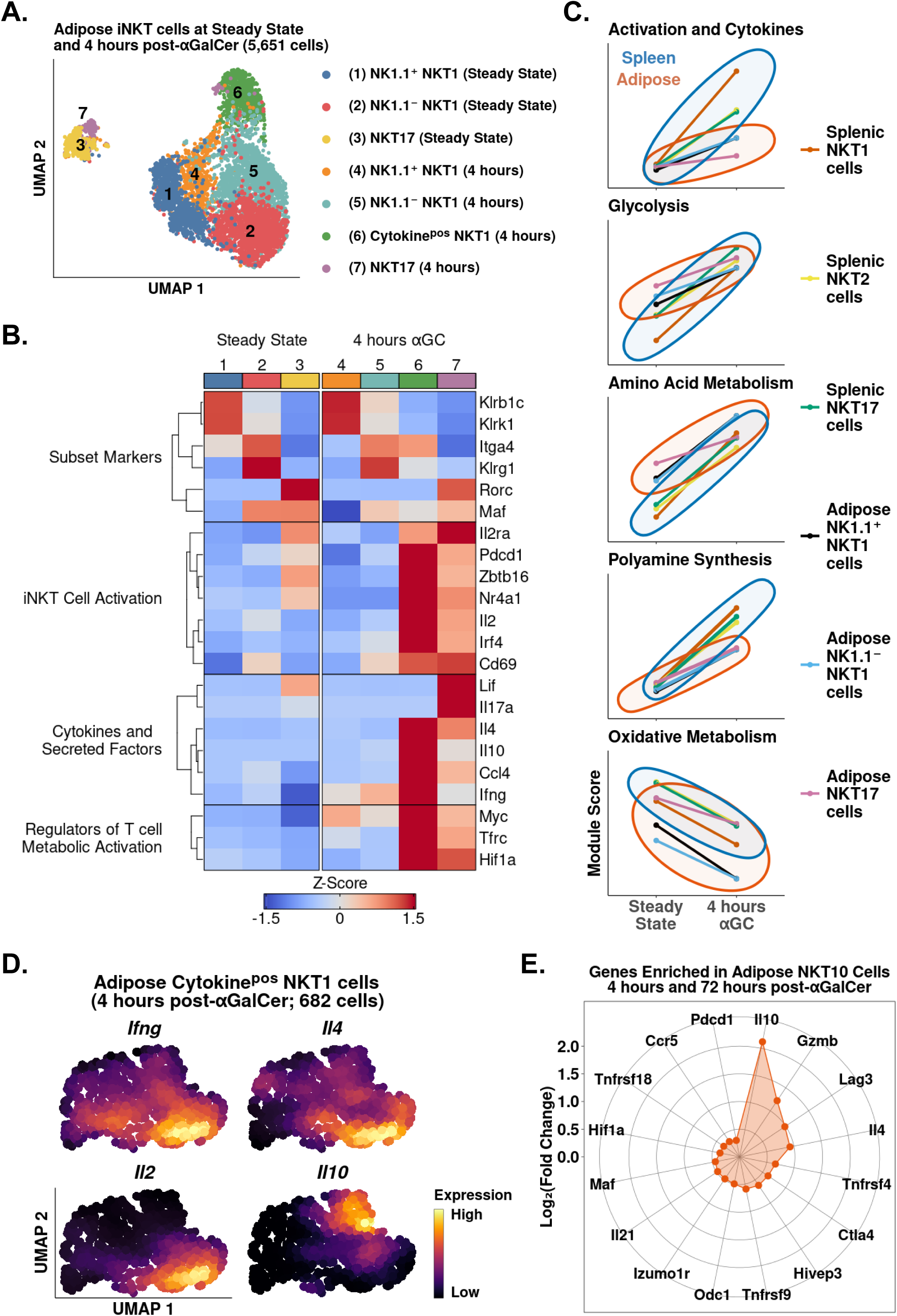
scRNA-Seq identifies transcriptional signatures of adipose iNKT cell subset activation. **A.** UMAP of murine adipose iNKT cells. **B.** Heatmap of averaged gene expression with hierarchical clustering in the data from Figure 4A. **C.** Line plots showing median expression of functional and metabolic module scores in adipose and splenic NKT1, NKT2 and NKT17 cell subsets. **D.** Density plots of gene expression in adipose cytokine^pos^ NKT1 cells 4 hours post-αGalCer (Cluster 6, Figure 4A). **E.** Radar chart showing Log_2_(Fold Change) values of genes enriched in Il10^pos^ adipose iNKT cells versus Il10^neg^ adipose iNKT cells 4 hours and 72 hours post-αGalCer.

Analysis of cytokine production among adipose iNKT cells revealed that *Il10* was only expressed by NKT1 cells (Figure 4B). Since cytokine^pos^ adipose NKT1 cells (Cluster 6) lacked or had downregulated expression of *Klrb1c* (NK1.1) by 4 hours post-αGalCer (Figure 4B), we could not stratify cytokine production in adipose NKT1 cells using *Klrb1c*. Therefore, we performed unbiased fine clustering of cytokine^pos^ adipose NKT1 cells. We identified one population of cells co-expressing *Ifng*, *Il4* and *Il2*, and a second population expressing *Il10* (Figure 4D), suggesting that adipose NKT10 cells are a distinct population. Analysis of cytokine^pos^ adipose iNKT cells at 72 hours post-αGalCer, when expression of *Il10* was highest among adipose iNKT cells (Figure 3), identified some NKT10 cells co-expressing *Il10* with *Ifng*, *Il4* and *Il21*, and other NKT10 cells that co-expressed *Il10*, *Ifng* and *Gzmb*, suggesting that expanded NKT10 cells are heterogeneous (Supplemental Figure 3). To investigate the transcriptional profile of adipose NKT10 cells we performed global analysis of *Il10*^pos^ versus *Il10*^neg^ adipose iNKT cells at 4 hours and 72 hours post-αGalCer. We identified 207 genes enriched in adipose NKT10 cells (Table S5), including the Tr1 cell markers *Lag3*, *Ctla4*, *Pdcd1*, *Gzmb*, *Il21*, *Hif1a*, *Maf* and *Ccr5*^42,44,48,49^, suggesting that adipose NKT10 cells may be functionally similar to Tr1 cells. In summary, our analysis suggests that adipose iNKT cells primarily segregate by function after αGalCer, and that regulatory adipose NKT10 cells are a transcriptionally distinct population similar to Tr1 cells.

### Chronic activation of splenic iNKT cells induces an adipose-like phenotype and the emergence of Tr1 iNKT cells

Since adipose iNKT cells displayed blunted activation after αGalCer, and enrichment of Tr1 cell markers, we wondered whether these were conserved features of regulatory iNKT cell biology. To answer this question we repeatedly activated splenic iNKT cells, which induces IL-10 production^19^. We sequenced 5,433 αGalCer activated splenic iNKT cells, including 2,117 cells isolated 4 weeks after mice received one dose of αGalCer (Resting) and 3,316 cells isolated 4 hours after reactivation with a second dose of αGalCer (Reactivated; Figure 5A). Comparison of iNKT cells at steady state (no αGalCer) and resting (4 weeks post-αGalCer) revealed that resting iNKT cells displayed a greatly reduced response to restimulation with αGalCer, similar to the adipose iNKT cell response to one dose of αGalCer (Figure 5B). This indicates that blunted activation after αGalCer is a conserved feature of regulatory iNKT cells. Furthermore, gene expression analysis revealed that resting NKT cells were transcriptionally similar to adipose iNKT cells, displaying reduced *Ifng*, *Il4* and *Il2* expression, and increased expression of Tr1 cells markers such as *Gzmb*, *Il10*, *Maf*, *Il21*, *Ctla4*, *Hif1a*, *Ccr5* and *Lag3*, especially after αGalCer re-challenge (Figure 5C), suggesting that regulatory iNKT cells are similar to Tr1 cells. We also reanalyzed previously published microarray data of control and αGalCer-pretreated splenic iNKT cells (GSE47959)^19^ (Supplemental Figure 4) and identified a similar activation phenotype in αGalCer-pretreated splenic iNKT cells versus control iNKT cells after short term αGalCer stimulation. Interestingly, although resting iNKT cells expressed *Tox* (Figure 5C), we did not identify enrichment of other chronic activation markers, such as *Nr4a1* (Nur77) (Supplemental Figure 5, Table S6), indicating that prior exposure of splenic iNKT cells to antigen does not completely reproduce the phenotype of adipose iNKT cells, which may be exposed to chronic endogenous activation *in situ*.

**Figure 5:**
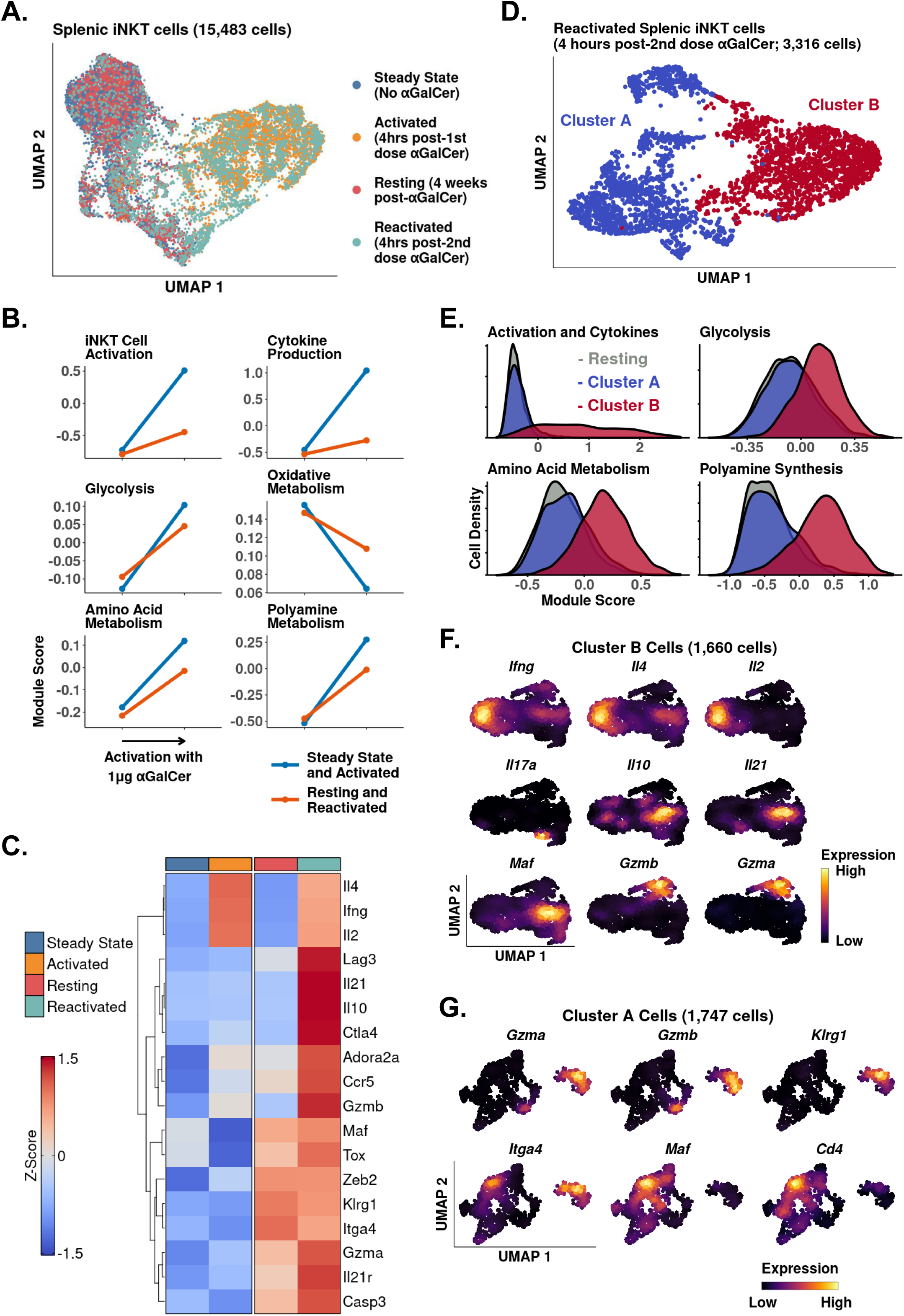
Chronic activation of splenic iNKT cells induces an adipose-like phenotype and the appearance of populations expressing Tr1 cell markers. **A.** UMAP of murine splenic iNKT cells. **B.** Line plots showing median expression of functional and metabolic module scores in the data from Figure 5A. **C.** UMAP of subclustered reactivated iNKT cells from Figure 5A. **D.** Histograms showing expression of functional and metabolic module scores in the data from Figure 5C. **E.** Heatmap of averaged gene expression with hierarchical clustering in the data from Figure 5A. **F.** Density plots of gene expression in iNKT cells from Cluster B cells from the data in Figure 5C. **G.** Density plots of gene expression in iNKT cells from Cluster A cells from the data in Figure 5C.

Since we had identified a distinct population of NKT10 cells in adipose tissue after αGalCer, we wondered whether we could also identify an NKT10 population among reactivated splenic iNKT cells. Fine clustering of reactivated splenic iNKT cells revealed two major populations, Cluster A and Cluster B (Figure 5D), which were differentiated by their response to reactivation. Only cluster B cells demonstrated significant activation, cytokine transcript expression, and metabolic reprogramming after the second dose of αGalCer (Figure 5E). Stratification of cluster B cells identified NKT1 cells expressing *Ifng*, *Il4* and *Il2*, NKT17 cells expressing *Il17a*, an NKT10 population expressing *Il10*, *Il21*, *Maf*, and intermediate *Ifng* and *Il4*, and a population of cells expressing *Gzmb*, *Gzma*, and intermediate *Ifng*, but not *Il4* (Figure 5F). In Cluster A, which was characterized by greatly blunted responsiveness to reactivation with αGalCer, there was a distinct population expressing *Gzmb*, *Gzma*, and the memory-like markers *Klrg1* and *Itga4* (Figure 5G)^8,10^. Moreover, although we could not stratify cells by *Il10* or *Il21* expression, we identified a population expressing *Maf*, *Itga4* and *Cd4*, which has been shown to be a marker of *Il10*^pos^ splenic iNKT cells after repeated αGalCer activation^19^. Therefore, this data suggests that repeated antigen exposure induces a regulatory phenotype in splenic iNKT cells, associated with the appearance of two novel memory-like populations expressing immunoregulatory cytokines and *Maf*, or cytotoxic markers and *Klrg1*.

### Memory-like cMAF^+^ and KLRG1^+^ iNKT cells are induced in the spleen following **α**GalCer challenge, and similar populations are constitutively present in adipose tissue

Having identified enrichment of iNKT cells expressing *Klrg1* and *Maf* among reactivated splenic iNKT cells, we wondered whether we could identify similar populations among resting iNKT cells. We also sought to characterize the transcriptional profile of these populations. Granular analysis of steady state and resting iNKT cells revealed distinct NKT1, NKT2 and NKT17 cell populations, demonstrating that iNKT cell subset diversity is preserved after antigen challenge (Figure 6A). Among resting iNKT cells, however, 2 new clusters emerged, expressing *Maf* or *Klrg1* (Figure 6A, 6B), matching our reactivated data. Flow cytometry of resting versus steady state splenic iNKT cells identified significantly more cMAF^+^ and KLRG1^+^ iNKT cells (Figure 6C, Figure 6D), demonstrating that cMAF^+^ and KLRG1^+^ iNKT cells are induced after antigen challenge. Gene expression analysis found that cMAF^+^ cells displayed enrichment of memory and exhaustion markers associated with memory precursor effector cells and T precursor exhausted cell populations, including *Slamf6*, *Tcf7*, *Ctla4*, *Lag3*, *Tox* and *Pdcd1* (Figure 6B)^12,25,50^. We also detected enrichment of *Zbtb16*, *Izumo1r* (FR4) and *Cd4*, and the Tr1 cell markers *Il27ra*, *Maf* and *Hif1a* (Figure 6B)^42,44,49^. By contrast, KLRG1^+^ cells showed enrichment of CD8^+^ effector T cell and NK cell markers such as *Gzmb*, *Gzma*, *Klrg1*, *Ncr1*, *Klrd1*, *Tbet* and *Klre1* (Figure 6B). KLRG1^+^ cells also expressed *Spry2* and *Cx3cr1* (Figure 6B), which were previously identified in KLRG1^+^ iNKT cells by Murray *et al*.^10^, and the transcription factor *Zeb2*. ZEB2 is a key transcription factor regulating terminal differentiation of KLRG1^+^ CD8^+^ effector cells^51^, suggesting that ZEB2 may be a candidate regulator of KLGR1^+^ iNKT cells. cMAF^+^ and KLRG1^+^ cells expressed *Itga4*, suggesting that these populations are memory-like or ‘trained’^10^.

**Figure 6:**
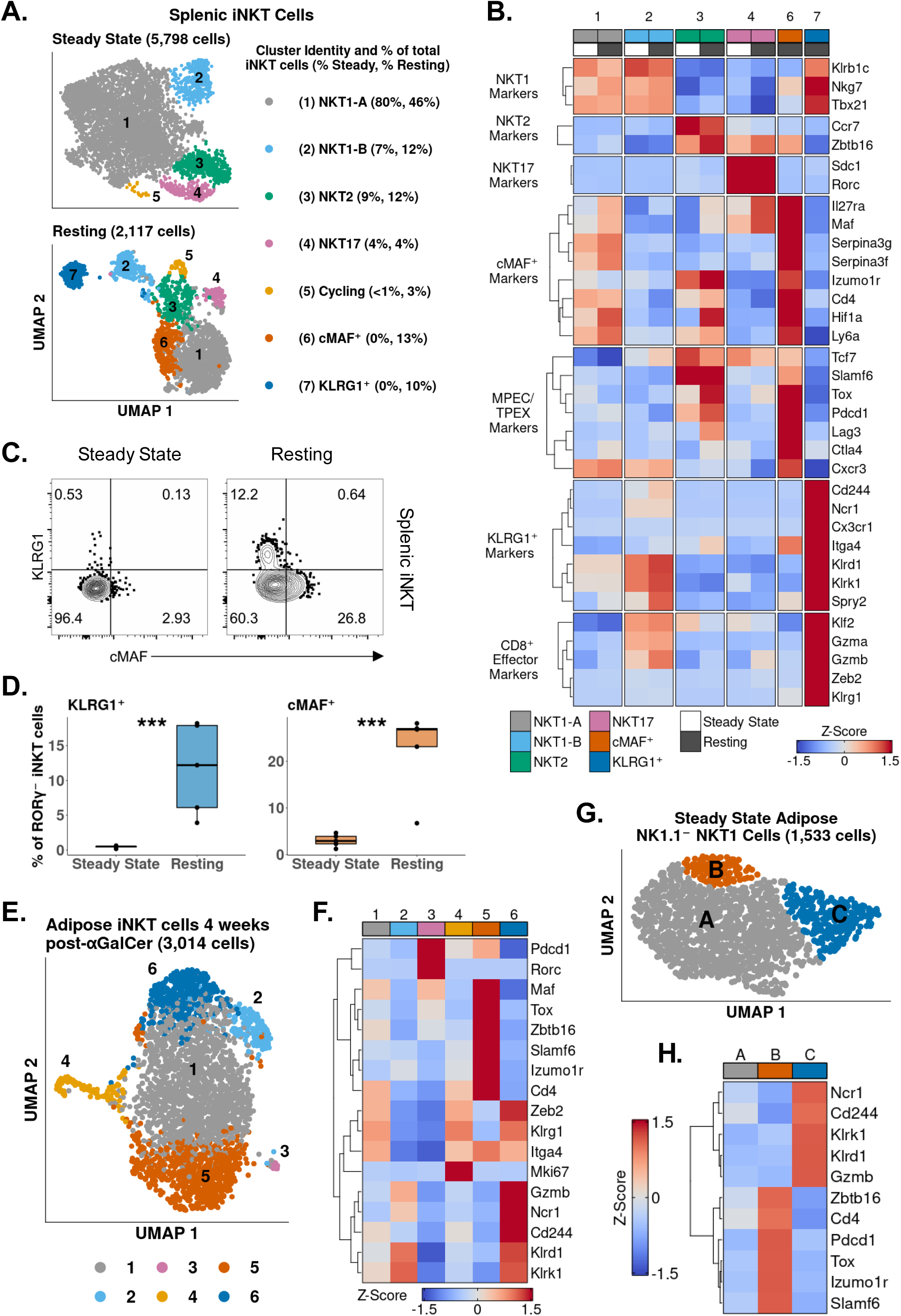
Memory-like cMAF^+^ and KLRG1^+^ iNKT cells are induced in the spleen following **α**GalCer challenge, and similar populations are constitutively present in adipose tissue. **A.** UMAP of murine splenic iNKT cells. **B.** Heatmap of averaged gene expression with hierarchical clustering in the data from Figure 6A. Both data sets were merged and normalized together, and cycling cells were excluded from the analysis. Memory precursor effector cell (MPEC), Precursor exhausted T cell (TPEX) **C.** Representative contour plots of RORγT^-^ splenic iNKT cells from untreated C57BL/6 mice (steady state) or C57BL/6 mice treated with a single dose of 4μg αGalCer four weeks prior to sacrifice (Resting). iNKT cells were defined as live, single CD19^-^ CD8^-^ CD11b^-^ CD3^low^ CD1d Tetramer^+^ cells and iNKT cells expressing RORγT were gated out. **D.** Boxplots showing percentages of cMAF^+^ or KLRG1^+^ iNKT cells as a percentage of RORγT^-^ splenic iNKT cells from steady state or resting C57BL/6 mice. N = 5 biological replicates from one experiment. Experiment performed once. Student’s unpaired t-test. Asterisks denote significance, * p < 0.05; ** p <0.01; *** p < 0.001. Horizontal lines represent median values. **E.** UMAP of murine adipose iNKT cells 4 weeks post-αGalCer. **F.** Heatmap of averaged gene expression with hierarchical clustering in the data in the data from Figure 6E. **G**. UMAP of murine steady state adipose NK1.1-NKT1 cells. H. Heatmap of averaged gene expression with hierarchical clustering in the data in the data from Figure 6G.

Having identified memory-like splenic cMAF^+^ and KLRG1^+^ iNKT cell populations that appear after αGalCer challenge, we wondered whether analogous populations were also present in adipose tissue. scRNA-Seq of 3,014 adipose iNKT cells from mice 4 weeks post-αGalCer (resting) revealed 6 clusters (Figure 6E). Strikingly, we detected cMAF^+^ (Cluster 5) and KLRG1^+^ (Cluster 6) populations (Figure 6F), indicating conserved enrichment of these two populations after αGalCer challenge across different tissues. Since adipose iNKT cells constitutively expressed markers of antigen experience and chronic activation, we also wondered whether adipose cMAF^+^ and KLRG1^+^ iNKT cells were present at steady state. Analysis of steady state adipose iNKT cells identified three distinct subpopulations (Figure 6G). All subpopulations expressed *Itga4*, *Maf* and *Klrg1* (Supplemental Figure 6), however, one population (Cluster B) expressed other cMAF+ cell markers such as *Izumo1r*, *Tox*, and *Slamf6* (Figure 6H), and another population (Cluster C) expressed KLRG1^+^ associated markers, including *Gzmb*, *Cd244* and *Klrk1* (Figure 6H). Overall, this data indicates that cMAF^+^ and KLRG1^+^ cell populations are induced by antigen experience in the spleen, and are constitutively present in adipose tissue.

### Identification of a conserved cMAF-associated and NKT_FH_-like transcriptional state in NKT10 cells

Having identified transcriptional signatures of regulatory iNKT cells in adipose tissue and after serial antigen activation, we next sought to describe shared transcriptional features of these different NKT10 cell populations. Gene expression analysis identified 110 genes enriched among splenic NKT10 cells (Table S7), including *Ctla4*, *Pdcd1*, *Lag3*, *Il21*, *Maf*, *Hif1a* and *Ccr5* (Figure 7A), all of which were already identified in adipose NKT10 cells. We identified 39 genes conserved across NKT10 cells from both tissues (Figure 7B), including Tr1 cell markers, *Tgfb1*, and the tolerogenic factors *Slfn2* and *Vsir*^52,53^ (Figure 7B). We also found that splenic NKT10 cells expressed the adipose iNKT cell marker *Nfil3* (Figure 7A), which we previously linked to IL-10 production by regulatory adipose NKT10 cells^17,18,54^. However, gene regulatory network analysis of *Il10* with *Nfil3* and other transcription factors identified in NKT10 cells using GENIE3^55^ revealed that *Maf* demonstrated the greatest correlation with *Il10* in NKT10 cells (Figure 7E). Since cMAF is known to regulate IL-10 in other immune populations, such as Tr1 cells, B cells and macrophages^49,56,57^, our data suggests that *Maf* is a major regulator of NKT10 cells. Flow cytometry of splenic iNKT cells 4 weeks post-αGalCer confirmed that IL-10^+^ iNKT cells expressed cMAF, and cMAF^neg^ cells produced little IL-10 (Figure 7F, Figure 7G).

**Figure 7:**
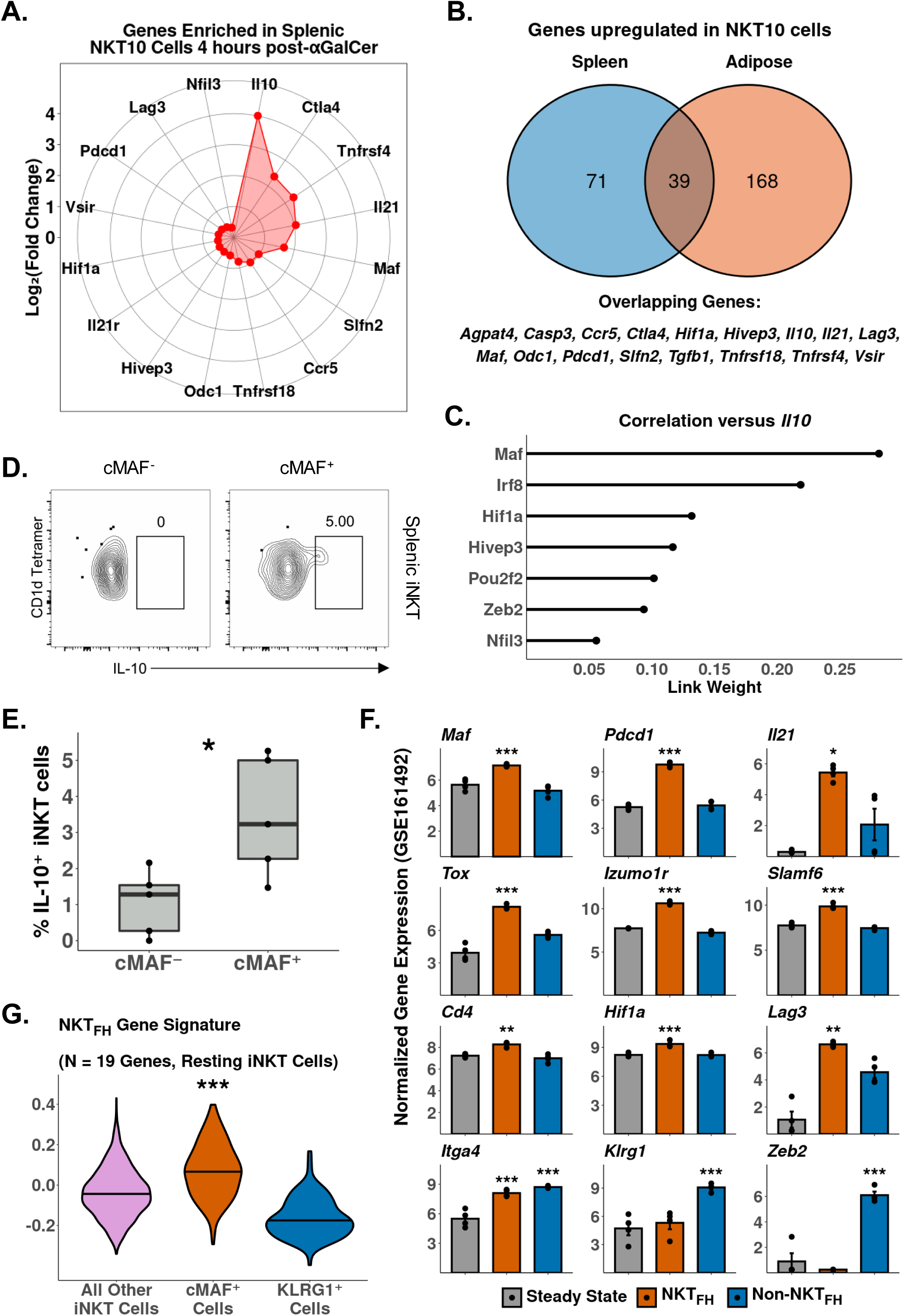
Identification of a conserved cMAF-associated and NKT_FH_- like transcriptional state in NKT10 cells. **A.** Radar chart showing Log_2_(Fold Change) values of genes enriched in cytokine producing *Il10*^pos^ iNKT cells versus *Il10*^neg^ splenic iNKT cells 4 hours post-αGalCer. **B.** Venn Diagram showing overlap of genes enriched in splenic (left) and adipose (right) *Il10*^pos^ iNKT cells after αGalCer (4 hours post-αGalCer for splenic iNKT cells, and 4 hours and 72 hours post-αGalCer for adipose iNKT cells). **C.** Lollipop plot showing correlation (Link Weight) of transcription factors versus *Il10* from GENIE3 analysis of NKT10 cells. **D.** Representative contour plots of IL-10 expression in cMAF^-^ or cMAF^+^ splenic iNKT cells after 4 hours of stimulation with 50ng PMA and 1µg Ionomycin (*ex vivo*). iNKT cells were isolated from C57Bl/6 mice treated with a single dose of 4μg αGalCer 4 weeks prior to sacrifice. iNKT cells were defined as live, single CD19^-^ CD8^-^ CD11b^-^ CD3^low^ CD1d Tetramer^+^ cells. **E.** Boxplots showing the percentage of total cMAF^-^ or cMAF^+^ iNKT cells expressing IL-10. N = 5 biological replicates from one experiment. Experiment performed once. Students t-test. Asterisks denote significance, * P < 0.05; ** P <0.01; *** P < 0.001. Horizontal lines represent median values. **F.** Barplots showing bulk RNA-Seq gene expression values in untreated total splenic iNKT cells (steady state) and splenic NKT_FH_ or Non-NKT_FH_ cells (6 days post-αGalCer). Reanalyzed from GSE161492. Asterisks indicate significantly increased expression versus all other populations or versus steady state alone (*Itga4* only). * P < 0.05; ** P <0.01; *** P < 0.001. **G.** Violin plots showing expression of NKT_FH_ Gene Signature module scoring in Resting Splenic iNKT cells (Figure 6A, bottom panel), with NKT1-A, NKT1-B, NKT2, NKT17 and Cycling cell clusters pooled together (All other iNKT cells) and compared versus cMAF^+^ and KLRG1^+^ iNKT cell clusters. Horizontal lines represent median values.

We next sought to compare the transcriptional signature of NKT10/cMAF^+^ cells against other memory-like iNKT cell populations. Interestingly, reanalysis of published bulk iNKT cell RNA-Seq data 6 days post-αGalCer (GSE161492)^10^ revealed that NKT_FH_ cells but not steady state or non-NKT_FH_ cells expressed NKT10/cMAF^+^ markers, including *Maf*, *Cd4*, *Tox*, *Izumo1r*, *Slamf6*, *Hif1a* and *Lag3* (Figure 7F, Table S7), and we found co-enrichment of NKT10/cMAF^+^ and NKT_FH_ markers in published microarray data comparing αGalCer-pretreated versus steady state splenic iNKT cells (GSE47959) (Supplemental Figure 4B, Supplemental Figure 7, Table S8). Module scoring of our resting scRNA-Seq data demonstrated that NKT10/cMAF^+^ cells but not KLRG1^+^ cells are enriched for NKT_FH_ cell gene signatures (Figure 6G). Overall, this suggests that NKT10/cMAF^+^ cells are transcriptionally similar to NKT_FH_ cells, and these two memory-like iNKT cell populations may phenotypically and functionally overlap.

## Discussion

iNKT cells and other innate T cells, such as γδ T cells and MAIT cells, rapidly activate following antigen or cytokine stimulus and produce potent cytokine responses. In this study we characterized the transcriptional programs underpinning the iNKT cell response to antigen, revealing an initial phase of rapid cytokine production, *Zbtb16* upregulation and metabolic remodeling, which was followed by second phase of proliferation, further remodeling of metabolic programs, and acquisition of features associated with memory-like iNKT cells. This transcriptional framework was highly conserved across iNKT cells from different tissues and species, and between functionally distinct iNKT cell subsets, including NKT1, NKT2 and NKT17 cells. Interestingly, we identified many similarities with adaptive T cell activation, including early induction of biosynthesis and aerobic glycolysis gene signatures^58^, although in keeping with the poised phenotype of iNKT cells this occurred over hours instead of days, in concert with innate-like cytokine production, and metabolic activation and cytokine production largely diminished by 3 days post-activation.

We sequenced 48,813 iNKT cells, the largest number of iNKT cells analyzed by scRNA-Seq to date, which allowed us to explore iNKT cell heterogeneity in unprecedented detail. We found that NKT2 and NKT17 cells, despite sharing many features of activation with NKT1 cells, were more enriched for genes associated with oxidative metabolism after activation, and production of NKT2 and NKT17 cell cytokines was more dependent on oxidative phosphorylation compared to NKT1 cell cytokines. Recent work by Raynor *et al.* (2020) has shown that NKT2 and NKT17 cells are more enriched for oxidative metabolism than NKT1 cells during thymic development, and downregulation of oxidative metabolism was required for establishment of an NKT1 cell phenotype^59^. Here we show that that enrichment of oxidative metabolism persists in splenic NKT2 and NKT17 cells after thymic development and defines their function, and we also identified enrichment of oxidative gene signatures in adipose NKT17 cells. Interestingly, we have shown that γδ17 cells display enrichment of oxidative metabolic signatures^39^, suggesting that enrichment of oxidative metabolism may be a conserved feature of IL-17 producing innate T cells.

Our study found that regulatory iNKT cell populations exhibited a blunted response to αGalCer, coupled with increased expression of early exhaustion markers. This agrees with previous studies demonstrating long term anergy in iNKT cells treated with αGalCer^7,19^ and enrichment of regulatory iNKT cells after prior αGalCer challenge^19^. Interestingly, we found that adipose iNKT cells, which are constitutively enriched for NKT10 cells, showed evidence of chronic endogenous activation, and repeated αGalCer challenge induced an adipose-like phenotype in splenic iNKT cells. Thus, chronic activation can promote a type of iNKT cell anergy, characterized by enrichment of regulatory iNKT cells. Chronic activation is known to promote exhaustion and anergy in adaptive T cells^45,46^, and a recent study by Vorkas *et al*. (2021) found that chronically activated human MAIT cells upregulate expression of FOXP3 and early exhaustion markers such as PD-1 and LAG-3^60^. Functionally, induction of regulatory and/or anergic innate T cells following chronic activation could serve as a protective mechanism to prevent over-activation of innate T cells, and limit immunopathogenicity. This could be relevant in autoimmunity, and it is notable that chronic activation of iNKT cells with αGalCer has been shown to ameliorate experimental autoimmune encephalomyelitis^61,62^.

Characterization of regulatory iNKT cells revealed two distinct memory-like iNKT cell populations, a cMAF^+^ population and a KLRG1^+^ population. KLRG1^+^ and cMAF^+^ iNKT cells were enriched in the spleen after prior antigen exposure, and were constitutively present in adipose tissue regardless of activation status. KLRG1^+^ iNKT cells showed enrichment of gene signatures associated with effector CD8^+^ T cells and NK cells, including cytotoxicity markers, and expressed the transcription factor ZEB2, which may regulate KLRG1^+^ iNKT cells. cMAF^+^ iNKT cells were transcriptionally similar to precursor exhausted T cells and NKT_FH_ cells, and were enriched for IL-10 producing NKT10 cells. We also reanalyzed bulk RNA-Seq data recently published by Murray *et al*. and found that NKT_FH_ cells were transcriptionally similar to cMAF^+^ iNKT cells, and expressed the transcription factor *Maf*, while non-NKT_FH_ cells were enriched for markers of KLRG1^+^ iNKT cells. Moreover, reanalysis of microarray data published by Sag *et al.* revealed co-enrichment of cMAF^+^, NKT_FH_ and KLRG1^+^ iNKT cell markers among splenic iNKT cells after prior antigen exposure. This suggests that antigen experience promotes the emergence of two distinct lineages of memory-like iNKT cells, an effector-like KLRG1^+^ population and an NKT_FH_-like population that expresses cMAF, and can produce IL-10. Interestingly, regulatory T_FH_ cells that produce IL-10 have been described^63^, suggesting that cMAF^+^ iNKT cells could represent a similar phenotype among iNKT cells.

We have shown that E4BP4 (*Nfil3*) rather than FOXP3 regulates production of IL-10 by adipose iNKT cells^17,18^, and we identified *Nfil3* expression among adipose and splenic NKT10 cells. However, most adipose iNKT cells express E4BP4 and do not express IL-10, suggesting that other factors must regulate IL-10. Furthermore, E4BP4 has been shown to bind an intronic region of the IL-10 promoter^54^, suggesting that other promoter-associated transcription factors are likely required to effectively induce IL-10 production in iNKT cells. Here, we implicate the AP-1 family transcription factor cMAF as a candidate regulator of NKT10 cells, correlating with published literature identifying a role for cMAF in regulation of IL-10 production by macrophages, B cells and adaptive T cells, including Tr1 cells^42,49,56,57^. cMAF is also expressed by NKT17 cells, and is required for their development^64^, as well as for γδ17 cell development^65^. However, although NKT17 cells can produce IL-10 after *in vitro* expansion^5^, we and others did not identify co-expression of IL-10 and IL-17 among *in viv*o or *ex vivo* iNKT cells^19^. Interestingly, RORγT has been demonstrated to suppress production of IL-10 in Th17 cells^66^, suggesting that it could perform a similar role in iNKT cells. We found that NKT10 cells co-expressed *Ifng*, *Il4*, and *Il21* with *Il10*, and lacked *Rorc* expression, indicating that NKT10 cells are more similar to NKT1 cells and NKT_FH_ cells than NKT17 cells. Notably, Murray *et al*. showed that most NKT_FH_ cells arise from T-bet^+^ NKT1 cells; given the transcriptional similarity of NKT10 and NKT_FH_ cells, it is possible NKT10 cells also arise from an NKT1 cell lineage. We also noted that NKT10 cells were transcriptionally similar to Tr1 cells, which are regulatory but lack expression of FOXP3, and are known to express cMAF^40–44,49^. Therefore, our data suggests that NKT10 cells may be similar to Tr1 cells^40,42,43^. Tr1 cells are heterogeneous, and we found that expanded adipose NKT10 cells were heterogeneous, with some cells expressing *Il4* and *Il21*, and other cells expressing *Gzmb*.

Understanding the factors controlling the phenotype and generation of regulatory iNKT cells has a several potential applications. For example, NKT10 cells have been shown to emerge in tumors, where they can promote tumor growth and regulatory T cell function^67^. However, our work suggests that NKT10 cells are capable of co-expressing *Ifng* and *Il10*. Skewing regulatory iNKT cells towards a less regulatory phenotype by enhancing IFNγ and inhibiting IL-10 might have a therapeutic benefit. We have shown that regulatory adipose iNKT cells can induce weight loss and suppress adipose tissue inflammation^68^, and are enriched in human adipose tissue^69^, suggesting that modulation of adipose iNKT cells might be beneficial during obesity and metabolic syndrome. Insights into the factors and transcriptional signatures governing iNKT cell activation are also potentially relevant to other innate T cell populations, as γδ T cells and MAIT cells respond similarly after activation^60,70^, and the transcriptomic resource that we present here may assist other studies investigating these innate T cell responses.

## Methods

### Animals

Male C57BL6 mice were purchased from Jackson Laboratory or Harlan Laboratory. BALB/C mice were purchased from Harlan Laboratory. Male and female BALB/C mice were bred under specific-pathogen-free facilities at Trinity College Dublin. Mice were used in experiments at 6-14 weeks of age. All animal work was approved and conducted in compliance with the Trinity College Dublin University Ethics Committee and the Health Products Regulatory Authority Ireland, and the Institutional Animal Care and Use Committee guidelines of The Dana Farber Cancer Institute and Harvard Medical School.

### *In vivo* stimulations

αGalCer (Avanti Polar Lipids) was prepared by dissolving in sterile DMSO (Sigma) at a concentration of 1mg/mL. αGalCer was prepared for injection by dilution in sterile PBS (Sigma) and the final concentration of DMSO was adjusted to less than 10% vol/vol. Mice were injected IP with 1µg of αGalCer or PBS vehicle. Mice were sacrificed and tissues harvested after either 4 or 72 hours. For analysis of memory-like iNKT cells mice were injected IP with 4µg of αGalCer and sacrificed after 4 weeks, with some animals recieving an additional IP injection of 1µg of αGalCer 4 hours before sacrifice. All injections had a final volume of 100µL.

### Tissue processing

Adipose tissue was excised, minced with a razor, and digested in 1mg/ml Collagenase Type II (Worthington) in RPMI shaking for 25-30 minutes at 37°C. Digested cells were filtered through a 70μM nylon mesh and centrifuged at 15,000 rpm for 5-7 minutes to pellet the stromovascular fraction (SVF). Spleens and thymi were disrupted through a 70μM filter and pelleted. Red blood cells in the spleen and thymus were lysed with ACK Lysing Buffer (VWR) or RBC Lysis Buffer (Biolegend) prior to further analysis.

### *Ex vivo* stimulations and mitochondrial staining

Where indicated, thymocytes were cultured for 4 hours in the presence of PMA and Ionomycin (Cell Stimulation Cocktail, Biolegend) and Brefeldin A (Biolegend), and in the presence or absence of oligomycin (Sigma). Cultures were in complete RPMI media supplemented with L-glutamine, penicillin, streptomycin and 10% FBS (Thermo Fisher Scientific). For mitochondrial staining, thymocytes were cultured for 30 minutes in complete RPMI media containing TMRM (Invitrogen) and/or Mitotracker Green FM (ThermoFisher Scientific).

### Flow cytometry and cell sorting

All antibody staining of live cells was performed in phosphate buffered saline (Gibco) with 1-2% FBS. Single cell suspensions were incubated in Fc-receptor blocking antibody (Clone 93, Biolegend) during cell surface antigen staining. Dead cells were excluded with Fixable Viability dyes (UV, eFluor 780; Thermo Fisher Scientific) or Zombie Aqua (Biolegend). For intracellular antigen staining, cells were fixed with either True-Nuclear Transcription Factor Buffer Set (Biolegend) for 45 minutes at room temperature, Foxp3/Transcription Factor Fixation/Permeabilization kit (Thermo Fisher Scientific) or Cytofix/Cytoperm kit (BD Biosciences) for 30 minutes at room temperature. The following anti-mouse antibodies were obtained from Biolegend: anti-CD3 (17A2), anti-CD19 (6D5), anti-CD11b (M1/70), anti-CD45 (30-F11), anti-TCRβ (H57-597), anti-CD8a (53-6.7), anti-KLRG1 (2F1/KLRG1), anti-IL-17A (TC11-18H10.1), anti-IL-4 (11B11), anti-IFNγ (XMG1.2), anti-CD43 Activation Glycoform (1B11), anti-ICOS (C398.4A). The following anti-mouse antibodies were obtained from Thermo Fisher Scientific: anti-cMAF (sym0F1), anti-RORγT (B2D), anti-RORγT (AFKJS-9), anti-CD8a (53-6.7), anti-IL-10 (JES5-16E3), anti-IL-13 (eBio13A), anti-CD11b (M1/70), anti-F4/80 (BM8). The following anti-mouse antibodies were obtained from BD Biosciences: anti-IFNγ (XMG1.2). iNKT cells were identified as live, single lymphocytes binding to anti-TCRβ or anti-CD3 antibodies and αGalCer analog PBS57-loaded CD1d tetramer (NIH Tetramer Core Facility/Emory Vaccine Center). A “dump” channel with antibodies against CD19, or CD19 and CD11b or F4/80, or CD19, CD8a and CD11b or F4/80 was used to eliminate nonspecific staining. For staining of BALB/C thymic iNKT cell subsets NKT1 cells were gated as CD43-HG^-^ ICOS^-^ CD3^low^ cells, NKT2 cells were gated as CD43-HG^-^ ICOS^+^ CD3^high^ cells, and NKT17 cells were gated as CD43-HG^+^ cells. Samples were acquired using LSR Fortessa and FACS Canto II cytometers and sorted using a FACS Aria Fusion Cell Sorter. Flow Cytometry analysis and plots were created using FlowJo version 10.0.7r2.

### scRNASeq sequencing and data pre-processing

Single-cell RNA-seq was performed on single cell suspensions of sorted iNKT cells from the visceral adipose tissue and spleens of mice using the 10x Genomics platform. A total of thirty-five visceral adipose tissue deposits or five spleens were pooled for each sample. Nine biological samples were sequenced in three batches (Table S9). For two of the batches, adipose and/or splenic iNKT cell samples were first tagged by TotalSeq-A Mouse hashtag antibodies (Biolegend) and then pooled for sequencing. Cell suspensions were loaded onto a 10x Chromium Controller to generate single cell Gel Beads-in-emulsion (GEMS) and GEMs were processed to generate UMI-based libraries according to the 10X Genomics Chromium Single Cell 3’ protocol. Libraries were sequenced using a NextSeq 500 sequencer (Illumina). Raw BCL files were demultiplexed using Cell Ranger v3.0.2 mkfastq to generate fastq files with default parameter. Fastq files were aligned to the mm10 genome (v1.2.0) and feature reads were quantified simultaneously using Cell Ranger count for feature barcoding. The resulting filtered feature-barcode UMI count matrices containing quantification of gene expression and hashtag antibody binding were then utilized for downstream analysis.

### Downstream scRNA-Seq data analysis

48,813 cells murine iNKT cells expressing a minimum median of 1,567 genes per cell and 4,163 UMIs per cell were loaded from feature-barcode UMI count matrices using the Seurat v4.0.3 package^71^. scRNA-Seq data for steady state adipose iNKT cells and splenic iNKT cells at steady state and 4 hours post-αGalCer were previously uploaded to GSE142845^18^. Raw human iNKT cell scRNA-Seq data was downloaded from GSE128243^36^. Antibody hashtag data was demultiplexed using the Seurat HTODemux function with a positive quantile of 0.99 and centered log ratio transformation normalization. Cells positive for more than one antibody were removed from the analysis. Genes expressed in less than three cells were excluded from all analysis to prevent false positive identification of transcripts. Cells expressing less than 1% minimum or more than 12.5% maximum of mitochondrial genes as a % of total gene counts were considered to represent empty droplets or apoptotic/dead cells and were removed from the analysis. Cells were also filtered based on total UMI counts and total gene counts on a per sample basis to remove empty droplets, poor quality cells and doublets, with a minimum cutoff of at least 500 genes per cell across all samples. UMI counts were normalized using regularized negative binomial regression using sctransform v0.3.2^71^. Where indicated, cycle regression was performed by first normalizing UMI counts using sctransform, then performing cell cycle scoring using the CellCycleScoring() function and cell cycle gene lists provided with the Seurat package, and then re-normalizing raw RNA count data with sctransform and regression of computed cell cycle scores applied.

Dimensionality reduction was performed using principal component analysis (PCA) with n = 100 dimensions and 2000 or 3000 variable features, and an elbow plot was used to determine the number of PCA dimensions used as input for uniform manifold approximation and projection (UMAP)^72^. For collective analysis of cells from different batches, the harmony v1.0 package^73^ was used with default settings to remove batch effects, and batch-corrected harmony embeddings were used for UMAP. Outlier cells expressing genes associated with macrophages (ex. *Adgre1*, *Cd14*), B cells (ex. *Cd19*) or CD8^+^ T cells (ex. *Cd8a*) were also identified and these cells were removed prior to the final analysis. UMAP was performed using a minimum distance of 0.3 and a spreading factor of 1. Shared nearest neighbor (SNN) graphs were calculated using k = 20 nearest neighbors. Graph-based clustering was performed using the louvain algorithm. In some cases, overclustering was performed and clusters were manually collapsed, and/or the first two dimensions of the UMAP reduction were used as input for graph-based clustering instead of PCA or harmony embeddings.

Gene expression analysis was performed using the FindMarkers() or FindAllMarkers() Seurat functions and the wilcoxon rank sum test. All gene expression analysis was performed using log-normalized RNA counts. Gene set enrichment analysis (GSEA)^38^ was performed using FGSEA v1.17.0^37^ and clusterProfiler v3.99.2^74^ packages. KEGG^30^ pathway data was retrieved from the Molecular signatures database (MSigDB)^75^ using the msigdbr v7.4.1 package. Over-representation analysis was performed using g:Profiler^76^. Heatmaps were generated using the Complex Heatmap v2.7.13 and circlize v0.4.13 packages^77^. Module scores were calculated using the AddModuleScore() Seurat function with n = 10 control features. Density plots were produced using the Nebulosa v1.1.1 package^78^. Gene regulatory network analysis was performed using GENIE3 v1.14.0^55^ with n = 10 iterations and link weight values were averaged between replicate analyses. Other plots were created using egg v0.4.5, GGally v2.1.2, ggiraphExtra v0.3.0, ggpubr v0.4.0, pals v1.7, patchwork v1.1.1, tidyverse v1.3.1, tidymodels v0.1.3, and viridis v0.6.1.

### Bulk RNA-Seq and microarray analysis

Raw RNA-Seq count files were downloaded from GEO Repository GSE161492^10^. Microarray data was downloaded from GEO Repository GSE47959^19^. Raw CEL files were annotated against the Mouse430_2 Array (mouse4302.db) and Robust Multichip Average (RMA) normalized using the affy v1.7.0 R package. Discrete probes corresponded to the same gene were merged and values were averaged. Raw RNA-Seq counts were transformed to log_10_(counts per million) and normalized using trimmed mean of M-values (TMM) normalization with EdgeR using edgeR v3.33.7^79^. Genes with low read counts were filtered out using the edgeR filterByExpr() function. Testing for differential gene expression was performed with Limma-Voom using limma v3.47.16^80,81^ with standard settings and without a minimum fold change cut-off. Plots were generated using the same libraries as used for scRNA-Seq data plotting.

### Statistics

Pilot studies were performed to estimate required sample size for adequate power. Significance was determined by Student’s two-tailed t test with Holm–Bonferroni correction, or one-way ANOVA with post-hoc Tukey test, where indicated. Significance is presented as *P < 0.05, **P < 0.01, ***P < 0.001, with P > 0.05 considered non-significant. All statistical analyses were performed using rstatix v0.7.0 and ggpubr v0.4.0 R packages. All sequencing data analysis was performed using R 4.1.2 and RStudio Desktop v1.4.1712 on an Ubuntu 20.04 Linux GNU (64 bit) system.

## Acknowledgments

The authors thank the Brigham and Women’s Hospital Single Cell Genomics Core for assistance with sequencing and pre-processing of scRNA-seq data, the NIH Tetramer Core Facility for recombinant CD1d PBS57 tetramers, and A.T. Chicoine for cell sorting. This work was supported by NIH grants R01 AI134861, American Diabetes Association 1-16-JDF-061 and ERC Starting grant 14283 (Project ID: 679173) (to L.L), and R01 AI113046 (to M.B.B.). Cartoons were created using Biorender.

## Author Contributions

H.K, N.M.L and A.N.S performed experiments. H.K. and N.M.L analyzed the experiments. H.K and L.L conceived the study. H.K performed all bioinformatic data analysis, and wrote the manuscript. L.L and M.B.B contributed to the design and interpretation of experiments and bioinformatic data analysis, and co-funded the study. L.L., N.M.L, and M.B.B edited the manuscript. L.L. supervised the study.

## Data Availability

scRNA-seq data from this manuscript have been deposited in the Gene Expression Omnibus under accession code GSE190201.

## Supplemental Figures

**Supplemental Figure 1.**
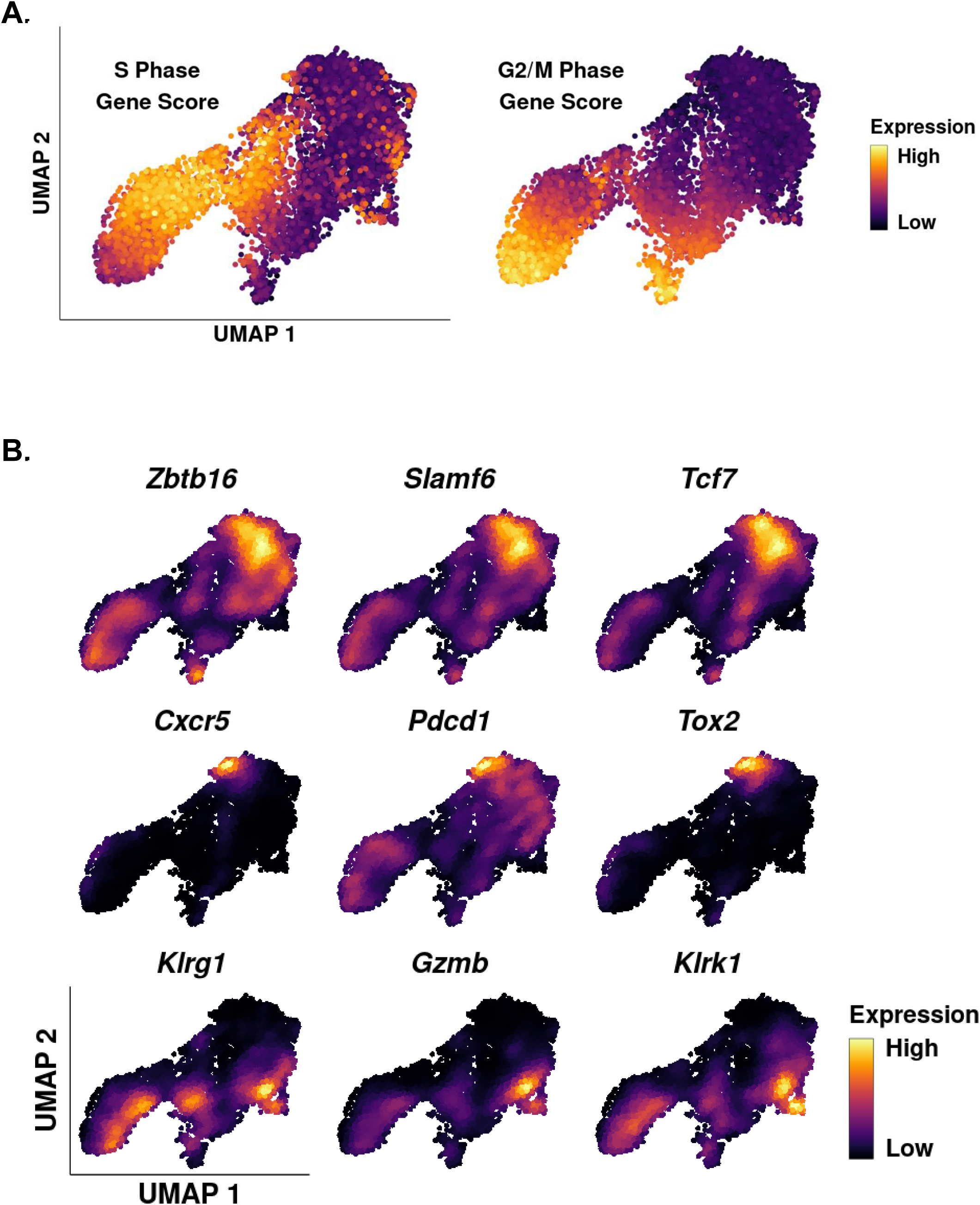
**A.** Expression plots of S Phase and G2/M phase proliferation gene scores in splenic iNKT cells 72 hours post-αGalCer. **B.** Density plots of gene expression in splenic iNKT cells 72 hours post-αGalCer.

**Supplemental Figure 2.**
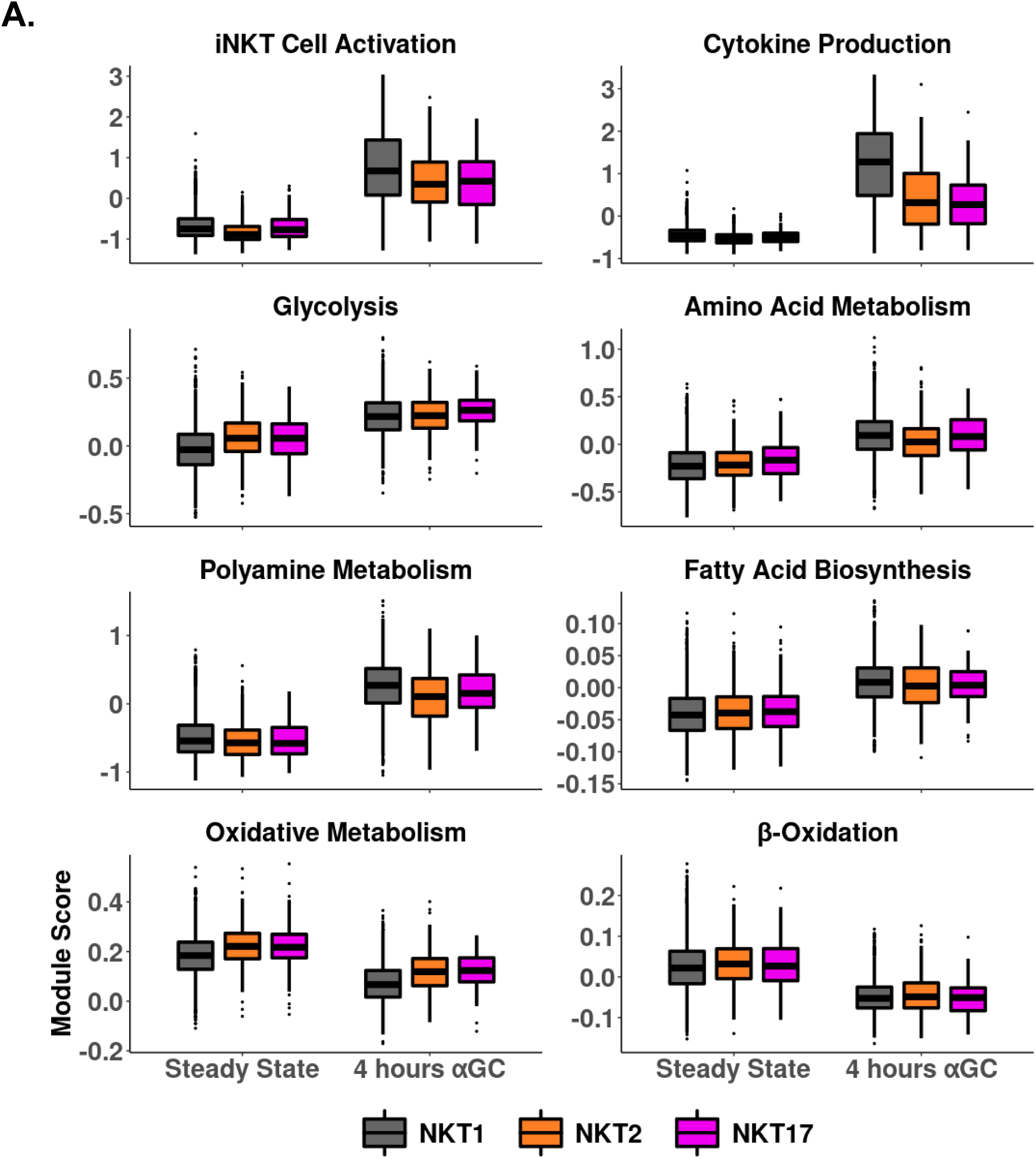
**A.** Boxplots of metabolic pathway gene module scores in scRNA-Seq data of NKT1, NKT2 and NKT17 cells at steady state and 4 hours post-αGalCer.

**Supplemental Figure 3.**
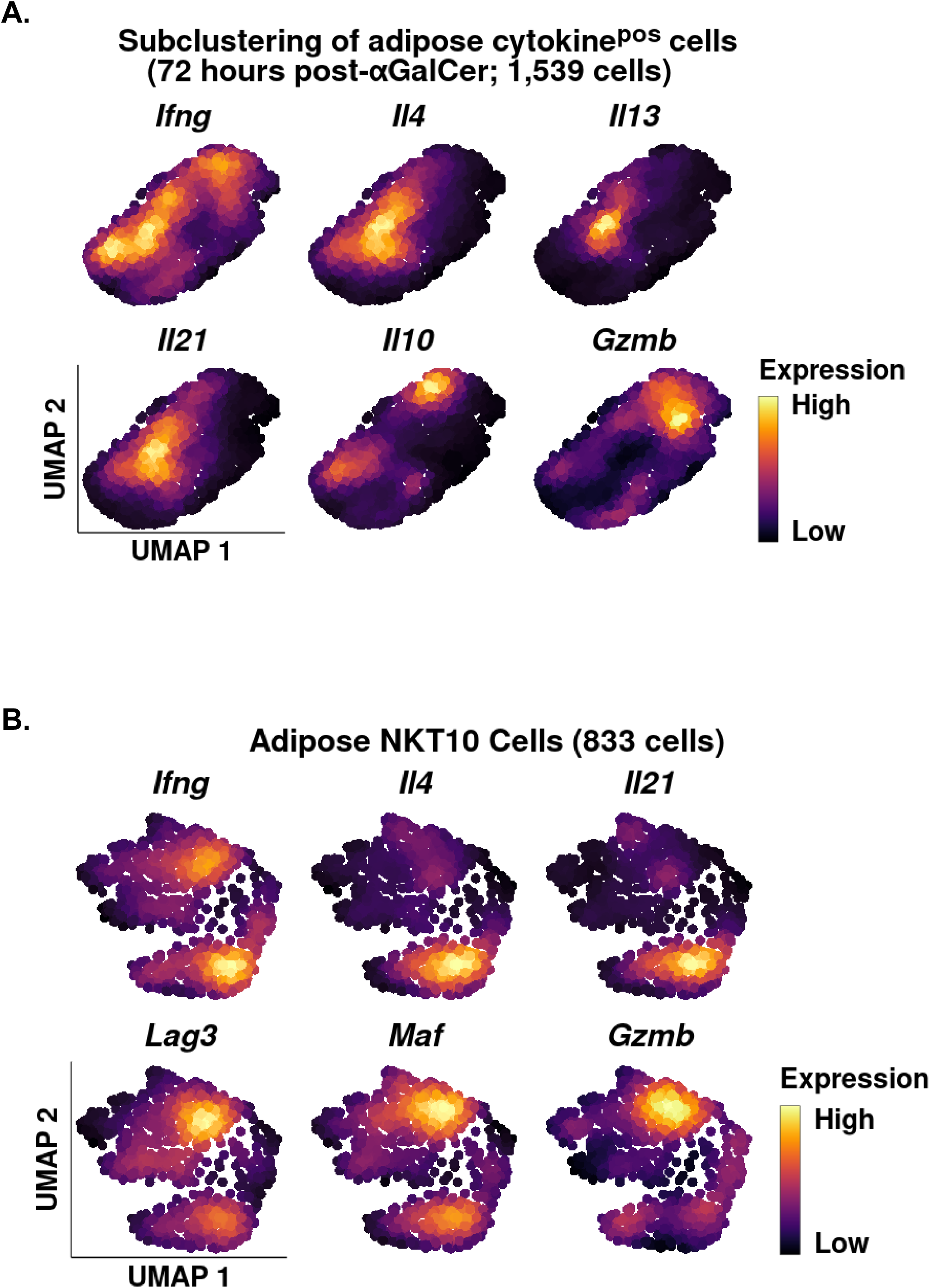
**A.** Density plots of gene expression in adipose cytokine^pos^ iNKT cells 72 hours post-αGalCer with cell cycle regression applied. **B.** Density plots of gene expression in adipose Il10^pos^ iNKT cells at 4 hours and 72 hours post-αGalCer with cell cycle regression applied.

**Supplemental Figure 4.**
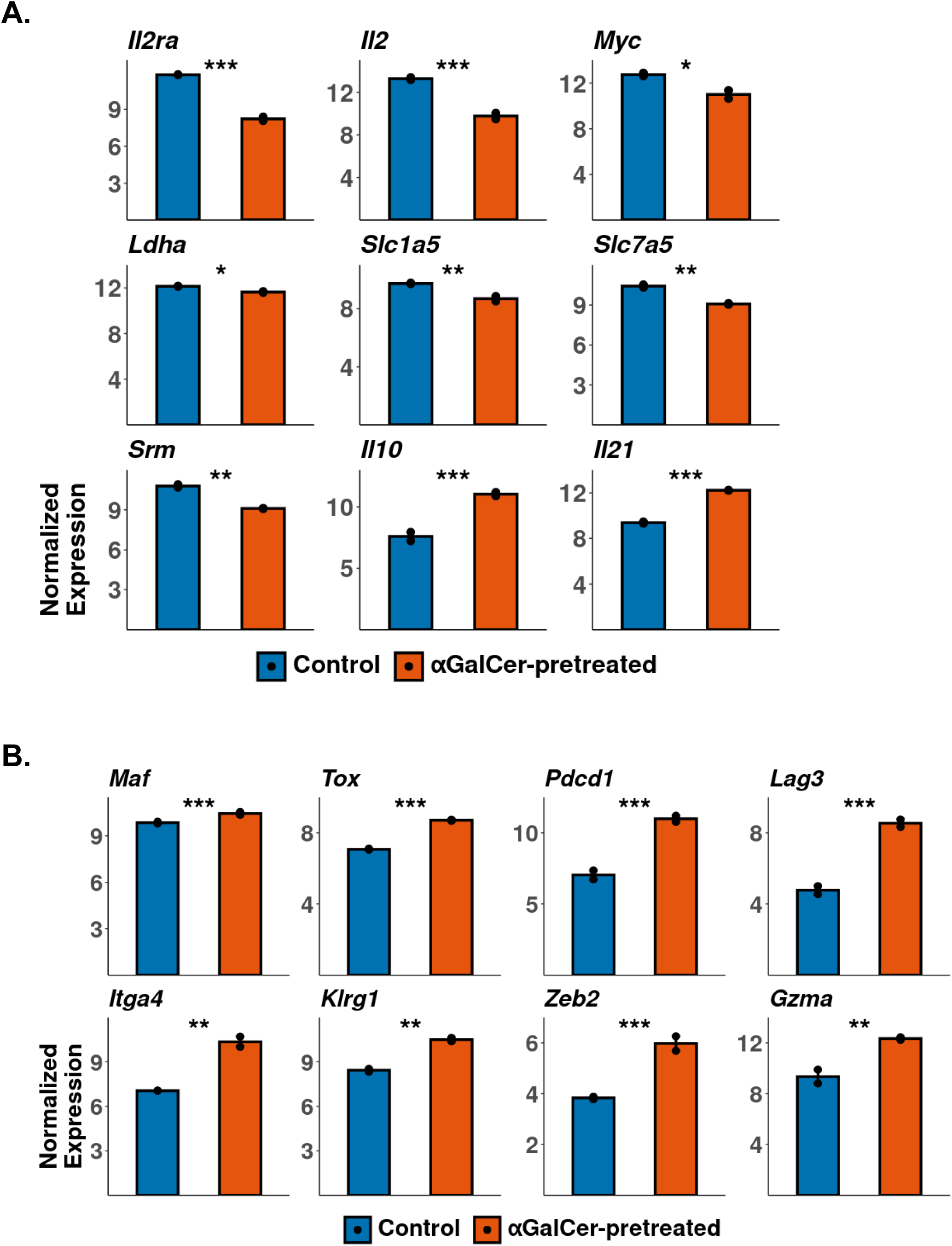
**A.** RMA-normalized gene expression in control and αGalCer-pretreated splenic iNKT cells both activated with 1μg αGalCer for 90 minutes *in vivo* prior to isolation. Reanalyzed from GSE47959. **B.** RMA-normalized microarray gene expression in control and αGalCer-pretreated splenic iNKT cells at rest. Reanalyzed from GSE47959.

**Supplemental Figure 5.**
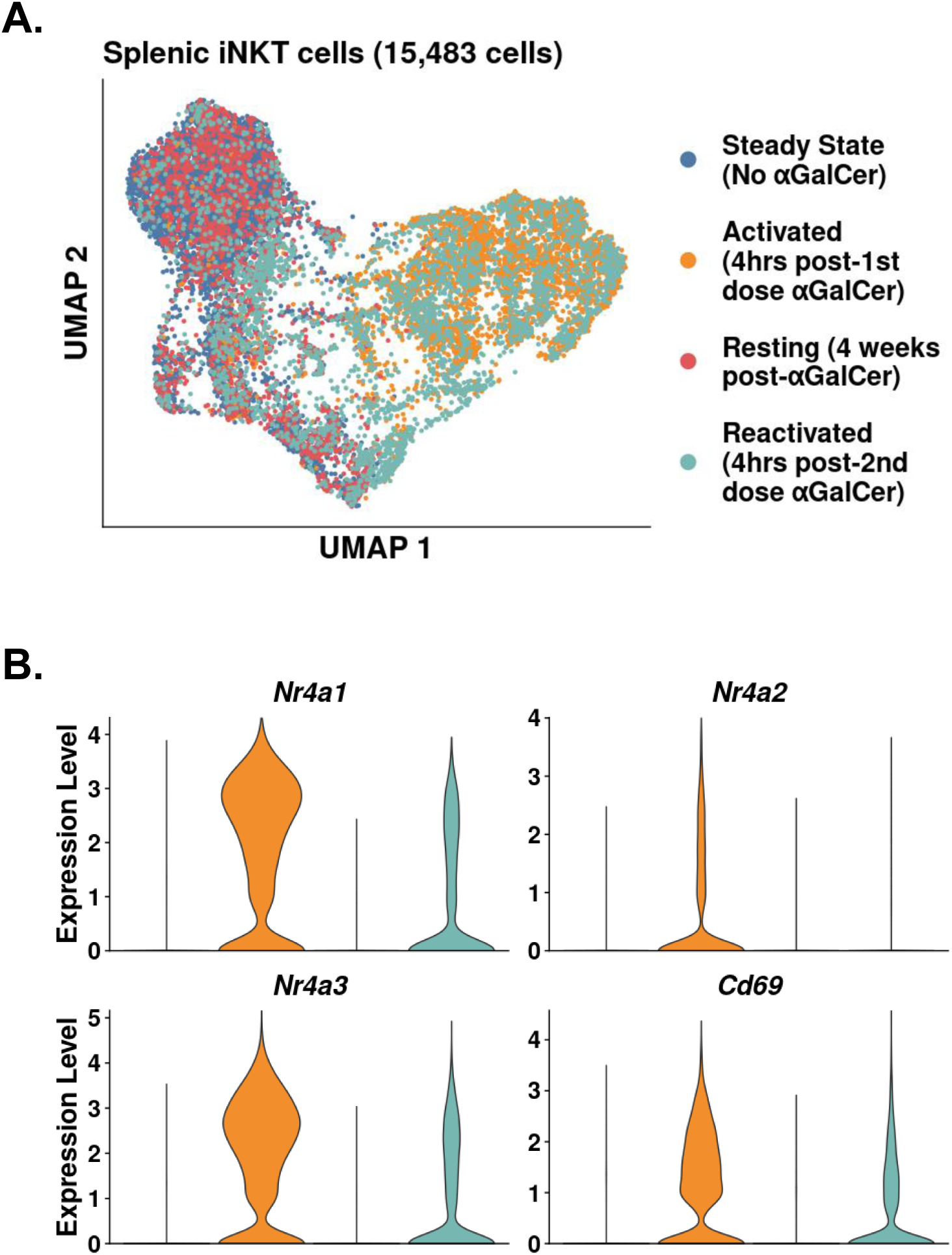
**A.** UMAP of murine splenic iNKT cells. **B.** Violin plots showing gene expression in the data from Supplemental Figure 5A.

**Supplemental Figure 6.**
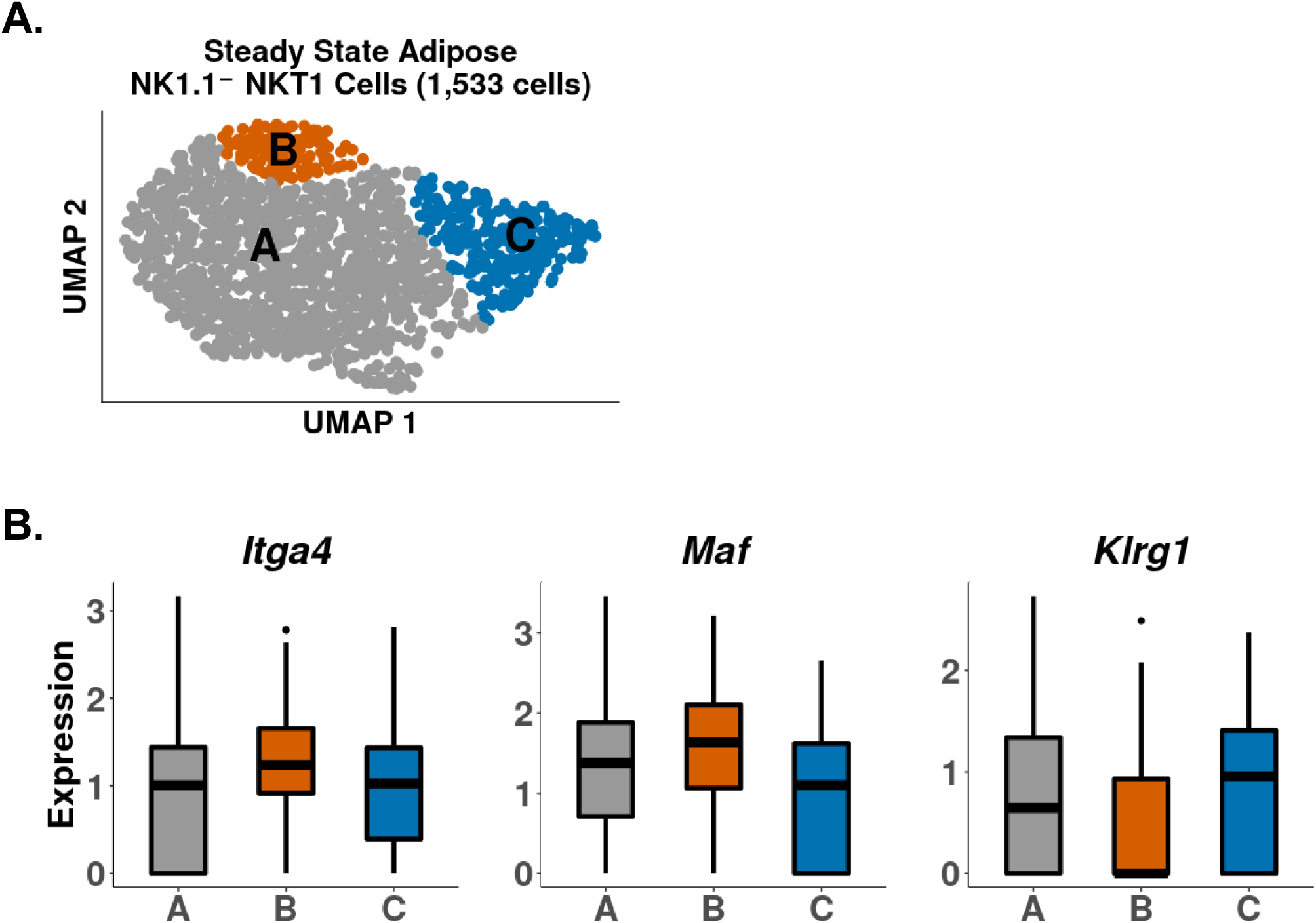
**A.** UMAP of murine steady state adipose NK1.1^-^ NKT1 cells. **B.** Boxplots showing gene expression in the data from Supplemental Figure 6A.

**Supplemental Figure 7.**
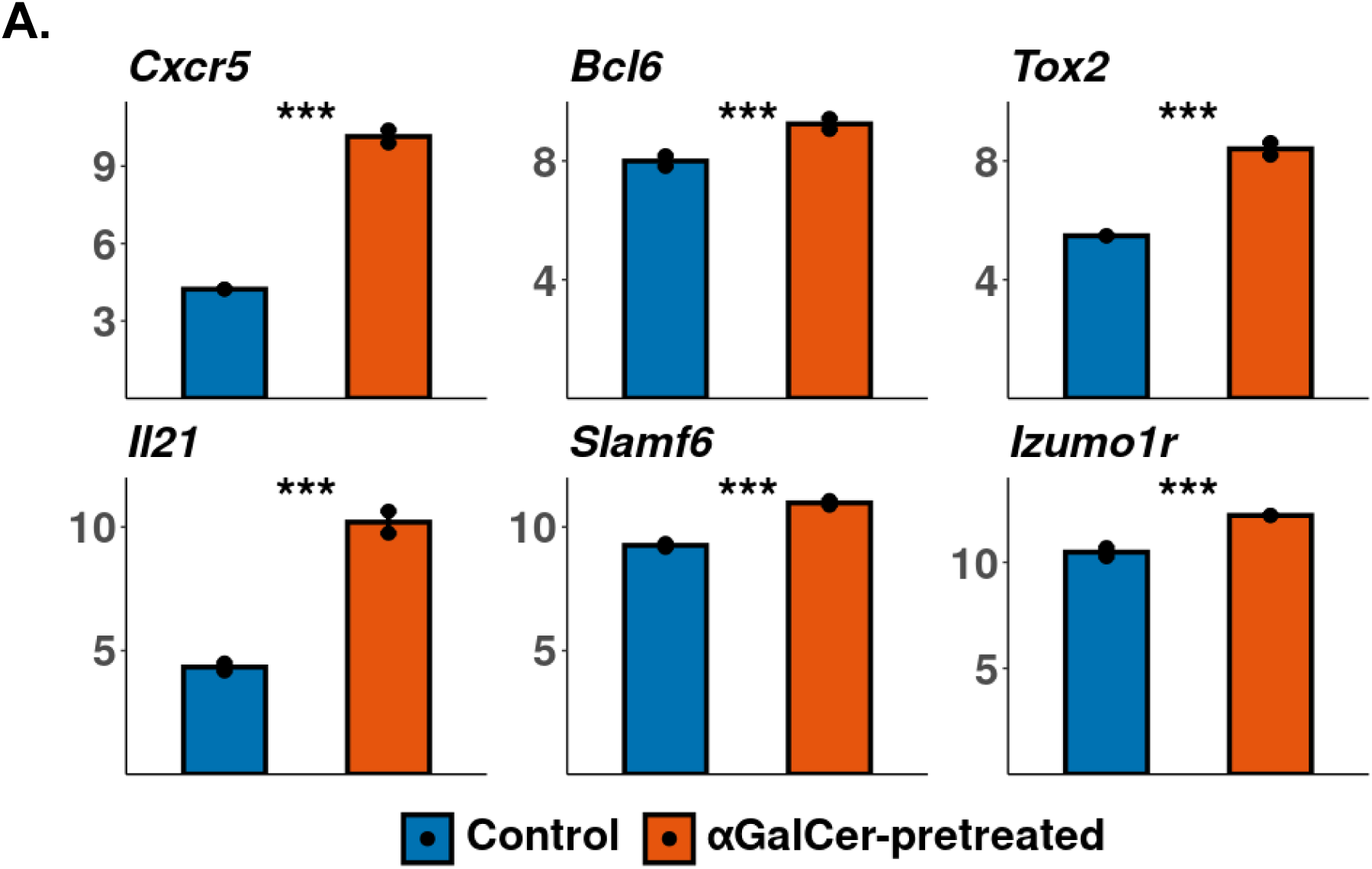
**A.** RMA-normalized microarray gene expression in control and αGalCer-pretreated splenic iNKT cells at rest. Reanalyzed from GSE47959.

## References

1. Kumar, B. V., Connors, T. & Farber, D. L. Human T cell development, localization, and function throughout life. Immunity 48, 202 (2018).

2. Chen, Y., Zander, R., Khatun, A., Schauder, D. M. & Cui, W. Transcriptional and Epigenetic Regulation of Effector and Memory CD8 T Cell Differentiation. Front. Immunol. 9, 2826 (2018).

3. Smyth, M. J. et al. Sequential activation of NKT cells and NK cells provides effective innate immunotherapy of cancer. J. Exp. Med. 201, 1973–1985 (2005).

4. Reilly, E. C. et al. Activated iNKT Cells Promote Memory CD8+ T Cell Differentiation during Viral Infection. PLoS One 7, e37991 (2012).

5. Cameron, G. & Godfrey, D. I. Differential surface phenotype and context-dependent reactivity of functionally diverse NKT cells. Immunol. Cell Biol. 96, 759–771 (2018).

6. Wilson, M. T. et al. The response of natural killer T cells to glycolipid antigens is characterized by surface receptor down-modulation and expansion. Proc. Natl. Acad. Sci. U. S. A. 100, 10913–10918 (2003).

7. Parekh, V. V. et al. Glycolipid antigen induces long-term natural killer T cell anergy in mice. J. Clin. Invest. 115, 2572 (2005).

8. Shimizu, K. et al. KLRG+ invariant natural killer T cells are long-lived effectors. Proc. Natl. Acad. Sci. U. S. A. 111, 12474–12479 (2014).

9. Chang, P.-P. P. et al. Identification of Bcl-6-dependent follicular helper NKT cells that provide cognate help for B cell responses. Nat. Immunol. 2011 131 13, 35–43 (2011).

10. Murray, M. P. et al. Transcriptome and chromatin landscape of iNKT cells are shaped by subset differentiation and antigen exposure. Nat. Commun. 12, 1–14 (2021).

11. Chen, Z. et al. Memory Follicular Helper Invariant NKT Cells Recognize Lipid Antigens on Memory B Cells and Elicit Antibody Recall Responses. J. Immunol. 200, 3117–3127 (2018).

12. Andreatta, M. et al. Interpretation of T cell states from single-cell transcriptomics data using reference atlases. Nat. Commun. 12, 1–19 (2021).

13. Engel, I. et al. Innate-like functions of natural killer T cell subsets result from highly divergent gene programs. Nat. Immunol. 17, 728–39 (2016).

14. Harsha Krovi, S., et al. Thymic iNKT single cell analyses unmask the common developmental program of mouse innate T cells. Nat. Commun. 11, 1–15 (2020).

15. Lee, Y. J. et al. Tissue-Specific Distribution of iNKT Cells Impacts Their Cytokine Response. Immunity 43, 566–578 (2015).

16. Baranek, T. et al. High Dimensional Single-Cell Analysis Reveals iNKT Cell Developmental Trajectories and Effector Fate Decision. Cell Rep. 32, (2020).

17. Lynch, L. et al. Regulatory iNKT cells lack expression of the transcription factor PLZF and control the homeostasis of Treg cells and macrophages in adipose tissue. Nat. Immunol. 16, 85–95 (2015).

18. LaMarche, N. M. et al. Distinct iNKT Cell Populations Use IFNγ or ER Stress-Induced IL-10 to Control Adipose Tissue Homeostasis. Cell Metab. 32, 243–258.e6 (2020).

19. Sag, D., Krause, P., Hedrick, C. C., Kronenberg, M. & Wingender, G. IL-10–producing NKT10 cells are a distinct regulatory invariant NKT cell subset. J. Clin. Invest. 124, 3725–3740 (2014).

20. Kovalovsky, D. et al. The BTB-zinc finger transcriptional regulator PLZF controls the development of invariant natural killer T cell effector functions. Nat. Immunol. 9, 1055–1064 (2008).

21. Stradner, M. H., Cheung, K. P., Lasorella, A., Goldrath, A. W. & D’Cruz, L. M. Id2 regulates hyporesponsive invariant natural killer T cells. Immunol. Cell Biol. 94, 640–645 (2016).

22. Marchingo, J. M., Sinclair, L. V., Howden, A. J. & Cantrell, D. A. Quantitative analysis of how myc controls t cell proteomes and metabolic pathways during t cell activation. Elife 9, (2020).

23. Finlay, D. K. et al. PDK1 regulation of mTOR and hypoxia-inducible factor 1 integrate metabolism and migration of CD8+ T cells. J. Exp. Med. 209, 2441–2453 (2012).

24. Wang, Z. et al. Iron Drives T Helper Cell Pathogenicity by Promoting RNA-Binding Protein PCBP1-Mediated Proinflammatory Cytokine Production. Immunity 49, 80–92.e7 (2018).

25. Utzschneider, D. T. et al. T Cell Factor 1-Expressing Memory-like CD8+ T Cells Sustain the Immune Response to Chronic Viral Infections. Immunity 45, 415–427 (2016).

26. Cohen, N. R. et al. Shared and distinct transcriptional programs underlie the hybrid nature of iNKT cells. Nat. Immunol. 14, 90–9 (2013).

27. Andris, F. et al. The transcription factor c-Maf promotes the differentiation of follicular helper T cells. Front. Immunol. 8, (2017).

28. Rampuria, P. & Lang, M. L. CD1d-dependent expansion of NKT follicular helper cells in vivo and in vitro is a product of cellular proliferation and differentiation. Int. Immunol. 27, (2015).

29. Carlson, C. M. et al. Kruppel-like factor 2 regulates thymocyte and T-cell migration. Nature 442, 299–302 (2006).

30. Kanehisa, M., Furumichi, M., Sato, Y., Ishiguro-Watanabe, M. & Tanabe, M. KEGG: Integrating viruses and cellular organisms. Nucleic Acids Res. 49, D545–D551 (2021).

31. Carbon, S. et al. The Gene Ontology resource: Enriching a GOld mine. Nucleic Acids Res. 49, D325–D334 (2021).

32. Angiari, S. et al. Pharmacological Activation of Pyruvate Kinase M2 Inhibits CD4+ T Cell Pathogenicity and Suppresses Autoimmunity. Cell Metab. 31, 391–405.e8 (2020).

33. Wu, R. et al. De novo synthesis and salvage pathway coordinately regulate polyamine homeostasis and determine T cell proliferation and function. Sci. Adv. 6, 4275–4291 (2020).

34. Fu, S. et al. Immunometabolism regulates TCR recycling and iNKT cell functions. Sci. Signal. 12, eaau1788 (2019).

35. Kumar, A. et al. Enhanced oxidative phosphorylation in NKT cells is essential for their survival and function. Proc. Natl. Acad. Sci. 116, 201901376 (2019).

36. Zhou, L. et al. Single-Cell RNA-Seq Analysis Uncovers Distinct Functional Human NKT Cell Sub-Populations in Peripheral Blood. Front. Cell Dev. Biol. 8, 384 (2020).

37. Korotkevich, G. et al. Fast gene set enrichment analysis. bioRxiv 060012 (2016) doi:10.1101/060012.

38. Subramanian, A. et al. Gene set enrichment analysis: A knowledge-based approach for interpreting genome-wide expression profiles. Proc. Natl. Acad. Sci. U. S. A. 102, 15545–15550 (2005).

39. Lopes, N. et al. Distinct metabolic programs established in the thymus control effector functions of γδ T cell subsets in tumor microenvironments. Nat. Immunol. 22, 179–192 (2021).

40. Yan, H. et al. Primary Tr1 cells from metastatic melanoma eliminate tumor-promoting macrophages through granzyme B- and perforin-dependent mechanisms. Tumor Biol. 39, (2017).

41. Gruarin, P. et al. Eomesodermin controls a unique differentiation program in human IL-10 and IFN-γ coproducing regulatory T cells. Eur. J. Immunol. 49, 96–111 (2019).

42. Chihara, N., Madi, A., Karwacz, K., Awasthi, A. & Kuchroo, V. K. Differentiation and characterization of Tr1 cells. Curr. Protoc. Immunol. 2016, 3.27.1–3.27.10 (2016).

43. Alfen, J. S. et al. Intestinal IFN-γ–producing type 1 regulatory T cells coexpress CCR5 and programmed cell death protein 1 and downregulate IL-10 in the inflamed guts of patients with inflammatory bowel disease. J. Allergy Clin. Immunol. 142, 1537–1547.e8 (2018).

44. Mascanfroni, I. D. et al. Metabolic control of type 1 regulatory (Tr1) cell differentiation by AHR and HIF1-α. Nat. Med. 21, 638 (2015).

45. Liu, X. et al. Genome-wide analysis identifies NR4A1 as a key mediator of T cell dysfunction. Nat. 2019 5677749 567, 525–529 (2019).

46. Seo, H. et al. TOX and TOX2 transcription factors cooperate with NR4A transcription factors to impose CD8+ T cell exhaustion. Proc. Natl. Acad. Sci. U. S. A. 116, 12410–12415 (2019).

47. Kumar, A. et al. Nur77 controls tolerance induction, terminal differentiation, and effector functions in semi-invariant natural killer T cells. Proc. Natl. Acad. Sci. 117, 17156–17165 (2020).

48. Grossman, W. J. et al. Differential expression of granzymes A and B in human cytotoxic lymphocyte subsets and T regulatory cells. Blood 104, 2840–2848 (2004).

49. Apetoh, L. et al. The aryl hydrocarbon receptor interacts with c-Maf to promote the differentiation of type 1 regulatory T cells induced by IL-27. Nat. Immunol. 11, 854–861 (2010).

50. Jadhav, R. R. et al. Epigenetic signature of PD-1+ TCF1+ CD8 T cells that act as resource cells during chronic viral infection and respond to PD-1 blockade. Proc. Natl. Acad. Sci. U. S. A. 116, 14113–14118 (2019).

51. Omilusik, K. D. et al. Transcriptional repressor ZEB2 promotes terminal differentiation of CD8+ effector and memory T cell populations during infection. J. Exp. Med. 212, 2027–2039 (2015).

52. Berger, M. et al. An Slfn2 mutation causes lymphoid and myeloid immunodeficiency due to loss of immune cell quiescence. Nat. Immunol. 2010 114 11, 335–343 (2010).

53. ElTanbouly, M. A. et al. VISTA is a checkpoint regulator for naïve T cell quiescence and peripheral tolerance. Science (80-. ). 367, (2020).

54. Motomura, Y. et al. The transcription factor E4BP4 regulates the production of IL-10 and IL-13 in CD4+ T cells. Nat. Immunol. 12, 450–459 (2011).

55. Huynh-Thu, V. A., Irrthum, A., Wehenkel, L. & Geurts, P. Inferring Regulatory Networks from Expression Data Using Tree-Based Methods. PLoS One 5, e12776 (2010).

56. Liu, M. et al. Transcription factor c-Maf is essential for IL-10 gene expression in B cells. Scand. J. Immunol. 88, (2018).

57. Cao, S., Liu, J., Song, L. & Ma, X. The Protooncogene c-Maf Is an Essential Transcription Factor for IL-10 Gene Expression in Macrophages. J. Immunol. 174, 3484 (2005).

58. Hartmann, F. J. et al. Single-cell metabolic profiling of human cytotoxic T cells. Nat. Biotechnol. 39, 186 (2021).

59. Raynor, J. L. et al. Hippo/Mst signaling coordinates cellular quiescence with terminal maturation in iNKT cell development and fate decisions. J. Exp. Med. 217, (2020).

60. Vorkas, C. K. et al. Single cell transcriptional profiling reveals helper, effector, and regulatory MAIT cell populations enriched during homeostasis and activation. bioRxiv 2020.10.22.351262 (2020) doi:10.1101/2020.10.22.351262.

61. Jahng, A. W. et al. Activation of Natural Killer T Cells Potentiates or Prevents Experimental Autoimmune Encephalomyelitis. 194, (2001).

62. Singh, A. K. et al. Natural Killer T Cell Activation Protects Mice Against Experimental Autoimmune Encephalomyelitis. J. Exp. Med. 194, 1801–1811 (2001).

63. Almanan, M. et al. IL-10–producing Tfh cells accumulate with age and link inflammation with age-related immune suppression. Sci. Adv. 6, eabb0806 (2020).

64. Thapa, P. et al. The differentiation of ROR-γt expressing iNKT17 cells is orchestrated by Runx1. Sci. Rep. 7, 7018 (2017).

65. Zuberbuehler, M. K. et al. The transcription factor c-Maf is essential for the commitment of IL-17-producing γδ T cells. Nat. Immunol. 20, 73–85 (2019).

66. Sun, M. et al. RORγt Represses IL-10 Production in Th17 Cells To Maintain Their Pathogenicity in Inducing Intestinal Inflammation. J. Immunol. 202, 79–92 (2019).

67. Wang, Y. et al. Unique invariant natural killer T cells promote intestinal polyps by suppressing TH1 immunity and promoting regulatory T cells. Mucosal Immunol. 2018 111 11, 131–143 (2017).

68. Lynch, L. et al. iNKT Cells Induce FGF21 for Thermogenesis and Are Required for Maximal Weight Loss in GLP1 Therapy. Cell Metab. 24, 510–519 (2016).

69. Lynch, L. et al. Invariant NKT cells and CD1d+cells amass in human omentum and are depleted in patients with cancer and obesity. Eur. J. Immunol. 39, 1893–1901 (2009).

70. Godfrey, D. I., Uldrich, A. P., McCluskey, J., Rossjohn, J. & Moody, D. B. The burgeoning family of unconventional T cells. Nat. Immunol. 16, 1114–1123 (2015).

71. Hafemeister, C. & Satija, R. Normalization and variance stabilization of single-cell RNA-seq data using regularized negative binomial regression. Genome Biol. 20, 1–15 (2019).

72. McInnes, L., Healy, J. & Melville, J. UMAP: Uniform Manifold Approximation and Projection for Dimension Reduction. arXiv (2018).

73. Korsunsky, I. et al. Fast, sensitive and accurate integration of single-cell data with Harmony. Nat. Methods 16, 1289–1296 (2019).

74. Yu, G., Wang, L. G., Han, Y. & He, Q. Y. ClusterProfiler: An R package for comparing biological themes among gene clusters. Omi. A J. Integr. Biol. 16, 284–287 (2012).

75. Liberzon, A. et al. Molecular signatures database (MSigDB) 3.0. Bioinformatics 27, 1739–1740 (2011).

76. Raudvere, U., et al. G:Profiler: A web server for functional enrichment analysis and conversions of gene lists (2019 update). Nucleic Acids Res. 47, W191–W198 (2019).

77. Gu, Z., Eils, R. & Schlesner, M. Complex heatmaps reveal patterns and correlations in multidimensional genomic data. Bioinformatics 32, 2847–2849 (2016).

78. Alquicira-Hernandez, J. & Powell, J. E. Nebulosa recovers single-cell gene expression signals by kernel density estimation. Bioinformatics 37, 2485–2487 (2021).

79. Robinson, M. D., McCarthy, D. J. & Smyth, G. K. edgeR: A Bioconductor package for differential expression analysis of digital gene expression data. Bioinformatics 26, 139–140 (2009).

80. Smyth, G. K. limma: Linear Models for Microarray Data. Bioinforma. Comput. Biol. Solut. Using R Bioconductor 397–420 (2005) doi:10.1007/0-387-29362-0_23.

81. Law, C. W., Chen, Y., Shi, W. & Smyth, G. K. Voom: Precision weights unlock linear model analysis tools for RNA-seq read counts. Genome Biol. 15, 1–17 (2014).

